# Function-driven geometry directs human pilosebaceous unit development

**DOI:** 10.64898/2026.08.31.745265

**Authors:** Elias Farr, Efpraxia Kritikaki, Marta Anna Chroscik, Chloe Admane, Emily Graves, Catherine Tudor, Hon Man Chan, Jacqueline Boccacino, Joseph McWilliam, Fereshteh Torabi, Keerthi Priya Chakala, Daniela Basurto-Lozada, Tong Li, Aljes Binkevich, Alexander Predeus, Martin Prete, Maryna Panamarova, Diana Adao, Kim Evans, Kris Stewart, Lloyd Steele, Elena Winheim, Hudaa Gopee, Emily Stephenson, Minal Patel, Christine Hale, Laure Gambardella, Beibei Du-Harpur, Catherine Smith, David Horsfall, Vijay Baskar, Leopold Parts, David J. Adams, Maria Kasper, Aurelien Dugourd, Julio Saez Rodriguez, April Rose Foster, Muzlifah Haniffa

**Affiliations:** Wellcome Sanger Institute, Wellcome Genome Campus, UK; University of Cambridge, Cambridge, UK; EMBL-EBI, Wellcome Genome Campus, UK; Newcastle University, Department of Dermatology and NIHR Newcastle Biomedical Research Centre, Newcastle, UK; Guys and St Thomas’ NHS Foundation Trust, Guy’s Hospital, Great Maze Pond, London SE1 9RT; St John’s Institute of Dermatology, King’s College London, Great Maze Pond, London SE1 9RT, United Kingdom; Department of Cell and Molecular Biology, Karolinska Institutet, Stockholm, Sweden; University Hospitals Foundation Trust, Addenbrooke’s Hospital, Cambridge, UK; Cambridge Institute for Therapeutic Immunology and Infectious Diseases (CIITID), University of Cambridge, UK

## Abstract

Single-cell technologies have generated cell censuses of tissues, however, how tissue geometry reflects functional needs remains poorly characterized. The human pilosebaceous unit offers a tractable model, a prenatally-formed complex mini-organ combining hair and sebum production with a stem cell reservoir. Using histomorphology, spatial transcriptomics, and single-cell multiomics on the same human prenatal scalp skin samples (8–19 post-conception weeks), integrated and analyzed using machine learning approaches, we built a spatiotemporal map of pilosebaceous unit development. We demonstrate that epithelial-mesenchymal interactions coordinate cellular fate and organogenesis, using an *in vitro* hair-bearing skin organoid model to validate this tissue-patterning. In addition, we show sebaceous gland developmental programmes are overcome during tumor formation. Our large-scale multi-modal analysis provides a unique framework for understanding form and function of tissues with applications in tissue engineering and pathology.

## Introduction

The relationship between form and function has long been studied to understand why animals and plants acquire the shapes they have during morphogenesis, primarily relying on mathematical (*1*) and biophysical (*2*) principles. In biology, function often refers to a trait or structure that causally contributes to an organism’s fitness (*3*). In the context of organs, which are assemblies of several tissues, these functions can be shared or divided among the different constituent tissues of the organ. In skin, this principle is exemplified by the pilosebaceous unit, a highly abundant mini-organ that encompasses the hair follicle (HF), sebaceous gland, and arrector pili muscle. With approximately 5 million distributed throughout the human body, pilosebaceous units provide a uniquely accessible model for studying organ development (*4*, *5*), including stem cell biology (*6–9*) and regeneration (*10*, *11*). The pilosebaceous unit is involved in many aspects of skin health, including sensory response, temperature control, light protection, and skin barrier integrity, in addition to its role in skin regeneration through specialized stem cell reservoirs. Disruption of its function and homeostasis contributes to disorders such as alopecia areata (*12*, *13*) and acne (*14*), and numerous genetic disorders resulting in hypertrichosis, hypotrichosis, and atrichia (*15*). Emerging evidence of the pilosebaceous unit harboring pathogenic T cells in inflammatory skin disease (*16*, *17*) demonstrates its role in contributing to disease memory, even after clinical response. Other diseases, including hidradenitis suppurativa (*18*) and sebaceous gland tumors (*19*), are associated with dysregulation or reactivation of developmental signalling pathways (*20*) such as Notch and Wnt during disease (*21*).

The functions of the pilosebaceous unit are provided by conserved architecture across mammals (*22*), whereby the hair follicle is organized into multiple cell layers that assemble along its vertical and medio-lateral axes. On the medio-lateral axis, this includes the outer mesenchymal layer and several inner epithelial layers (hair shaft, inner and outer root sheath) (Fig. 1A). On the vertical axis, the HF spans from the hair canal right to the hair bulb deep in the dermis, where transit amplifying cells reside that ultimately produce the hair shaft (Fig. 1A). At the midpoint of the vertical axis resides the stem cell-rich bulge region of the HF, which provides the progenitors for HF regeneration (*23*), wound repair (*11*), and an anchor point for the arrector pili muscle formation (*24*). Above the bulge sits the sebaceous gland, which produces sebum through holocrine secretion.

**Fig. 1:**
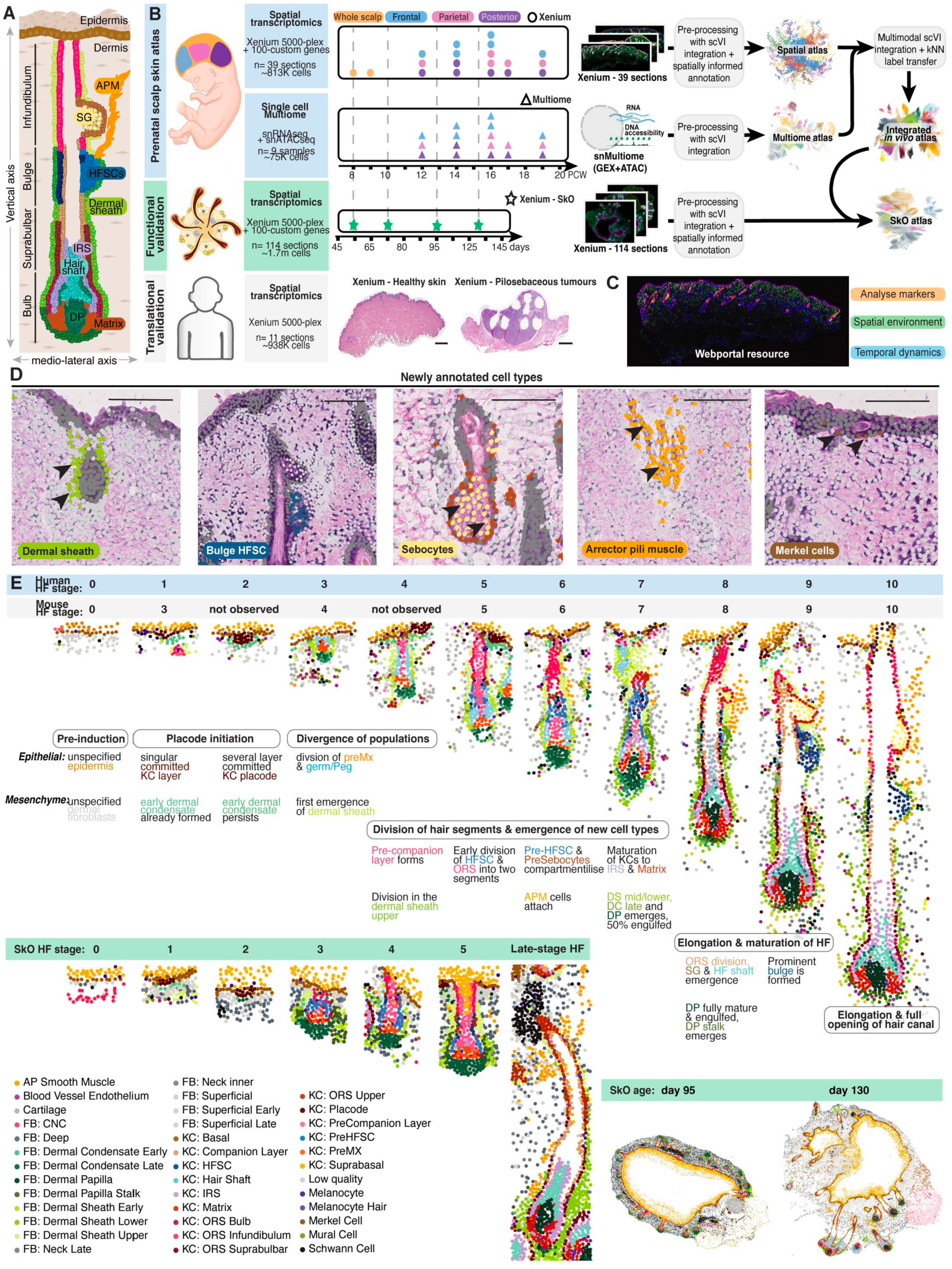
Spatial transcriptomics map all stages of human hair follicle development. A Cartoon highlighting pilosebaceous unit morphology B Dataset generation and analysis approach overview. Image-based spatial transcriptomics capturing 5000 genes plus 100 extra genes, together with Multiome (RNA and ATAC from the same cell), was generated from 3 scalp regions spanning PCW 8 to 19. In addition, functional validation in terms of images and hair-bearing SkOs was generated in the equivalent time span in days. C The dataset can be displayed and analyzed in a web portal, allowing analysis of markers, visualization of the spatial environment in the whole section, and the interrogation of spatial dynamics by visualizing genes across the different stages of hair development D Newly detected cell types from the in vivo prenatal skin atlas. Overlays of H&E staining with cell type annotation, showing the spatial location of newly captured cell types. E Cropped and aligned examples of HFs from Stage 0 to 10 (See Table 1) with developmental milestones and comparison to the iPSC-derived SkO HF development, bottom right: full sections of day 95 and day 130 organoids.

Tissue morphogenesis into a defined architecture is often tightly regulated, which requires local and global tissue coordination (*25*). While local coordination refers to interactions among neighboring cells within a tissue often directed by mechanical cues (*26–28*) and cell-cell signaling, global coordination refers to coordinated development between distant tissue compartments, such as the bulge and the bulb compartments in the pilosebaceous unit that together control hair growth.

Like many other organs, such as the kidney, lung, and several glands, HFs form prenatally in humans through reciprocal interactions of the epithelial (epidermis) and mesenchymal (dermis) compartments. HFs cycle through growth, resting, and shedding phases throughout postnatal life, which partially recapitulate development (*29*), but no new follicles are formed after birth. The earliest phase of HF formation results from epithelial-mesenchymal interactions and, in mice, is known to be driven by the Wnt (*30*, *31*), Shh (*32*, *33*), and Fgf (*34*) pathways. Local polarisation of epidermal cells (*35*) and recruitment of dermal fibroblasts from the placode and dermal condensate of the nascent HF (*36*, *37*). The role and impact of epithelial-mesenchymal interactions past the initiation, including the role of the dermal sheath, is less well studied.

Most of our knowledge on hair development comes from studies in rodents, and it is still unclear how much of those findings are directly translatable to humans. Functional differences between the species exist, including specialized rodent hair types that can develop in the same tissue space and are absent in humans, a shorter hair cycle, the onset of shaft production after birth in rodents that does not occur in humans, and hormone-related disorders (*38*) absent in mice. Furthermore, current classifications of HF morphogenesis focus on early murine development (*36*, *39*). There are currently no human HF developmental stages or cell states, and molecular characterization of many conserved stages, including dermal sheath, arrector pili muscle, and inner HF layer formation. On the cellular level, several stem cell populations (*7*, *40–42*) and markers including Lhx2 (*43*), Lgr5 (*41*), Lgr6 (*40*), Sox9 (*9*), and Nfatc1 (*44*) with distinct expression patterns along the vertical axis have been reported in mice but warrant validation to be present in humans. Lineage tracing studies in mice have revealed the importance of the progenitor cells’ position in the hair placode in determining their final cell fate (*6*). However, it is unclear if the cells are already committed in the placode and simply mature throughout HF formation or if there are additional signaling interactions between the epithelial progenitor cells and their mesenchymal and epithelial surroundings that determine their final fate.

The temporal dynamics of human HF development are challenging to study *in vivo* due to the scarcity and logistics of acquiring human prenatal skin. A recent advancement that can support this research is the complex *in vitro* iPSC-derived hair-bearing skin organoid (SkO) model, which recapitulates milestones in human skin development (*45*), including hair peg formation after 60 days and fully formed hair shafts after 130 days in culture (*46*). How well this hair formation follows the timeline of *in vitro* hair formation and whether there are functional differences between the skin organoid and prenatal skin has not been comprehensively described yet. However, as the model is derived from the neural crest, it only recapitulates facial and frontal scalp skin, and other limitations include failure to recapitulate adult cell states (47).

Studying human pilosebaceous unit development has relied on methods such as histopathological imaging (*37*) and single-cell RNA-sequencing (*47*, *48*), which provide either spatial or molecular information alone, making it difficult to relate cellular molecular states to tissue architecture during organogenesis. These complementary layers of information can be integrated by combining H&E staining, imaging-based spatial transcriptomics and single-cell multiomics, which measure the full transcriptome and epigenome from the same cell. Together, these approaches enable the study of local and global tissue development and determine how functional requirements shape the geometry of the pilosebaceous unit during morphogenesis.

Here, by generating a comprehensive dataset of human prenatal scalp skin using H&E staining, high-resolution spatial transcriptomics, and single-cell multiome (joint profiling of RNA and ATAC) from 8 to 19 PCW, we reveal the spatiotemporally coordinated development of the human pilosebaceous unit, including new marker genes for cell types previously indistinguishable by conventional imaging methods. We characterize: i) the local and global orchestration of the epithelial-mesenchymal interactions forming the hair placode and extending throughout HF development, ii) similarities between arrector pili muscle and cardiomyocyte differentiation, and iii) sebaceous gland formation, suggesting overriding of normal sebaceous gland developmental programmes in sebaceous gland tumors. Furthermore, we used the SkO model to validate and benchmark our *in vivo* prenatal HF developmental findings. Finally, we provide this dataset as a resource for the community in a web portal that allows interrogation of spatially resolved genes and cell types throughout the differentiation stages of HFs. Our findings establish the spatiotemporal logic of human pilosebaceous unit development and provide a developmental framework for understanding its biology in health and disease.

## Results

To investigate the development of human HFs in three regions (frontal, temporal, occipital) of prenatal scalp skin across induction and maturation (8-19 weeks post conception (PCW)), we leveraged a multi-omics approach combining histomorphological and transcriptome data from imaging-based spatial transcriptomics (Xenium, 5001 genes + 100 custom relevant to the pilosebaceous unit based on literature and scRNA-seq data) with haematoxylin and eosin (H&E) staining of the same tissue sections and single-nuclei RNA- and ATAC-sequencing (snMultiome) profiling of the same tissue blocks (Fig. 1B). Cell segmentation was aided by antibody staining of the plasma membrane, nuclear and cytoplasmic proteins. We used the platform manufacturer’s segmentation protocol, which showed similar performance when benchmarked against other segmentation methods (Fig. S1a).

After preprocessing, scVI (*49*, *50*) integration of snMultiome and spatial transcriptomics datasets, and spatially informed annotation (Fig. S1, S2, Methods), the combined dataset comprised ∼900k high-quality cells – 800 from the spatial transcriptomics of 39 sections and ∼100k cells for the Multiome – generating the first comprehensive multi-modal human HF development dataset (Fig. 1B). This dataset can be browsed interactively in a webportal (Fig. 1C). With 519 median genes and 813 median counts per cell, the spatial data is richer than published human or mouse skin datasets (*51*, *52*) (Fig. S1b). Using this data, we spatially resolved cell types, tissue niches, and signalling events during HF differentiation (Fig. 1B, Methods). We leveraged the open chromatin data to predict the molecular regulation underpinning cell type differentiation during HF development (Fig. 1B, Methods). We observed that combining the spatial and dissociated single-cell data allows for greater resolution in deconvolving cell types than using suspension scRNA-seq alone (Fig. S3a,b), as we recently reported (*52*).

We describe the transcriptomes and cellular location of developing cell types, including those of the pilosebaceous unit, some of them for the first time in humans, such as bulge hair follicle stem cells (HFSC), sebocytes, arrector pili muscle cells, dermal sheath, and rare cell types such as Merkel cells (Fig. 1D, S3a,b). The increased cell type resolution is conferred by a combination of factors, including not having to dissociate skin, which can damage vulnerable cell types (Sebocytes), additional spatial information to differentiate similar cells beyond their transcriptome (arrector pili muscle, dermal sheath), and having a richer dataset with higher cell numbers (Merkel cells). Table S1 provides the marker genes for all cell types. Our dataset captured several immune cell types, including myeloid cells like macrophages, mast cells, dendritic cells, Langerhans cells, and granulocytes (Fig. S4a). Apart from Langerhans cells and dendritic cells that are located in the epidermis and upper HF as expected (Fig. 1D, S4b), most of the myeloid cells were distributed in the dermis, as observed in mice (*53*) (Fig. S4b). A fraction (∼3 %) of macrophages were in the dermal sheath (Fig. S4c), as previously described (*54*, *55*), where they have been posited to have a maintenance function. We observed B and T lymphocytes around vessels as expected (Fig. S4b).

We generated imaging-based spatial transcriptomics (Xenium, 5001 genes + 100 custom) with H&E staining from the SkO model to benchmark it against *in vivo* HF development and for functional validation. Five time points of culture were selected to describe skin development and match prenatal scalp sampling: day 34 (after the dermis and epidermis specification but no HF initiation), day 54 (neural crest-derived cell types differentiate), day 76 (HFs morphologically visible, equivalent 9-10 PCW), day 95 (intermediate stage of HF differentiation, equivalent 14 PCW), and day 130 (formation of “mature” HFs, equivalent 17-19 PCW) (Fig. 1B). The SkO xenium data (∼1.7 m Xenium cells, 114 sections, 329 median genes per cell, 444 median total counts) were processed and annotated with the same approach as the *in vivo* data, using the same cell-type labels as the scalp data where relevant (Methods). As expected, the SkO dataset largely recapitulates the cell-type composition of the prenatal *in vivo* dataset, with some differences such as off-target cartilage cell types (*45*) and the absence of immune cells (*45*) and lymphatic endothelial cells (*47*) (Fig. S3c).

To derive a holistic classification across all stages of HF development and perform consistent comparison of histomorphology simultaneously with spatial transcription patterns from the same HF, we cropped and aligned all HFs that were fully captured within the spatial data of the skin sections to the same axis. The HF stages captured ranged from the earliest epithelium and mesenchyme condensation to the fully established pilosebaceous unit (Fig. 1D, Methods). After ordering the stages by developmental age (Methods), we analysed a total of 137 individual HFs from prenatal scalp skin (Table S2).

We divided these HFs into 10 developmental stages based on morphology and molecular information, providing the most comprehensive classification of hair development so far (Fig. 1E). Stages 1-4 describe HF induction and formation of placode and dermal condensate, stages 4-6 describe the the first visible appearance of the pre-companion layer (PreCL)(Stage 4), segregation of outer root sheath (ORS) segments (Stage 5) and arrector pili muscle induction (Stage 6), while stages 7-10 describe the maturation of the pilosebaceous unit components, including the inner root sheath (Stage 7), dermal papilla (Stage 8), sebaceous gland (Stage 8), hair shaft (Stage 8) and bulge (Stage 9)(Fig. 1E). When compared with previous murine classifications (*36*, *39*), key differences and additions emerge at Stages 1 to 4, whereas Stages 5-10 are conserved. Key differences include murine stages 1 and 2 from Saxena *et al.*, characterized by placodes without clustered dermal condensate, which are not observed in our human prenatal scalp dataset and therefore excluded in the human classification, whereas additions include two new stages that do not correspond to any murine HF differentiation stages (Table 1). These are human HF Stage 2, which represents a human-specific placode stage characterized by placode downgrowth without division into matrix and ORS progenitors as observed in mice (*39*, *56*), and human HF Stage 4, representing the appearance of the PreCL, which was described in mice (*57*) but not included in the murine classifications (Fig. 1E).

In addition, we provide a comprehensive comparison between prenatal scalp skin and the iPSC-derived SkO (*45*). This analysis shows conservation of both the presence of and spatial relationship between keratinocyte and mesenchymal populations in the first 5 stages of hair development, including dermal sheath and arrector pili muscle, but there was divergence thereafter, with the SkO HFs not undergoing stages 6-10 (Fig. 1E). HFs in both prenatal skin and SkO develop over 8 weeks but lack the hair bulge and sebaceous glands on day 130 of culture, in line with missing stages 6-10 (Fig. 1E). We provide an interactive visualization of the histomorphology and molecular data of individual tissue sections as well as the full spectrum of HF stages in a webportal.

### Human Hair Placode and Dermal Condensate Form Concurrently

To understand the molecular dynamics of the human prenatal scalp epidermal placode, we interrogated the Xenium spatial transcriptomics data for marker gene expression characteristic of hair placodes and histomorphological epithelial thickening. We observed intermittently spaced regions of the epithelium that expressed *FGF20*, *EDAR,* and *WNT10B*, defining the emerging placode, without concurrent evidence of epithelial thickening in the H&E section at 9 PCW (Fig. S5a,b). This finding puts the earliest evidence of placode formation in the human scalp one week earlier than previously reported (*48*) and demonstrates commitment to placode fate and expression of established markers without histologically discernible apical-basal elongation (*58*). Canonical placode thickening patterns, which involve apical-basal polarisation and nuclei elongation as visible in different regions of fetal skin (*48*), were observed from 12 PCW in humans; however, this was without expression of *SHH, a* critical driver of HF induction in mice (*33*) and expressed in human Stage 3 (Fig. S5c, Table S2). The percentage of placode cells formed per surface centimeter of prenatal scalp skin increased between 9 and 12 PCW (Fig. S5d,e). Placodes observed from 14 PCW tended to be next to a newly formed follicle (starting from Stage 4), and in 19 PCW, mature HFs are typically accompanied by two developing follicles, consistent with triplet HF formation previously described in mice (Fig. S5f)(*59*). New placodes also tend to colocate with existing HFs in the SkO at day 95 of culture (Fig. S5g). Collectively, this suggests that the first induction of HF differentiation happens at 8-9 PCW in the human prenatal scalp and is followed by one or several waves between 12-17 PCW.

Given hair regression from androgenetic alopecia preferentially affects the fronto-temporal regions of the male scalp (*13*, *60*), we assessed if this may result from placode abundance, growth rates, or differential gene expression established during prenatal scalp hair formation. However, no substantial qualitative or quantitative differences in placode abundance, HF growth rate, or gene expression were detected using Milo (*61*) or differential gene expression analysis between frontal, temporal, and occipital scalp regions (Fig. S5h-j).

We next sought to characterize human prenatal scalp placode differentiation in more detail. In murine HF development (*39*), initial commitment of the basal epithelial cells to placode formation at Stage 2 is directly followed by asymmetrical division within placodes into basal *SHH*+ pre-matrix (KC: PreMX) and apical *SOX9*+ KC: Germ/Peg cells at Stage 3, before the placode starts its downgrowth (*56*). Here we provide the first evidence that these cells are present in the hair pegs at Stage 3 of human HF development, which is after the HF has initiated downgrowth (Fig. 2A). At human HF Stage 2, which already shows invagination, placodes do not contain *SHH*+ (KC: PreMX) and *SOX9*+ (KC: Germ/Peg) cells (Fig. 2A, S5b). This observation is paralleled by a bifurcation in the inferred keratinocyte developmental trajectory starting from placode cells into *SHH*+ (KC: PreMX) and *SOX9*+ (KC: Germ/Peg) cells, which occurs at Stage 3 (Fig. S6a, Methods). Our findings demonstrate that human prenatal scalp PreMX and Germ/Peg lineage acquisition occurs at a later stage than shown in mice.

**Fig. 2:**
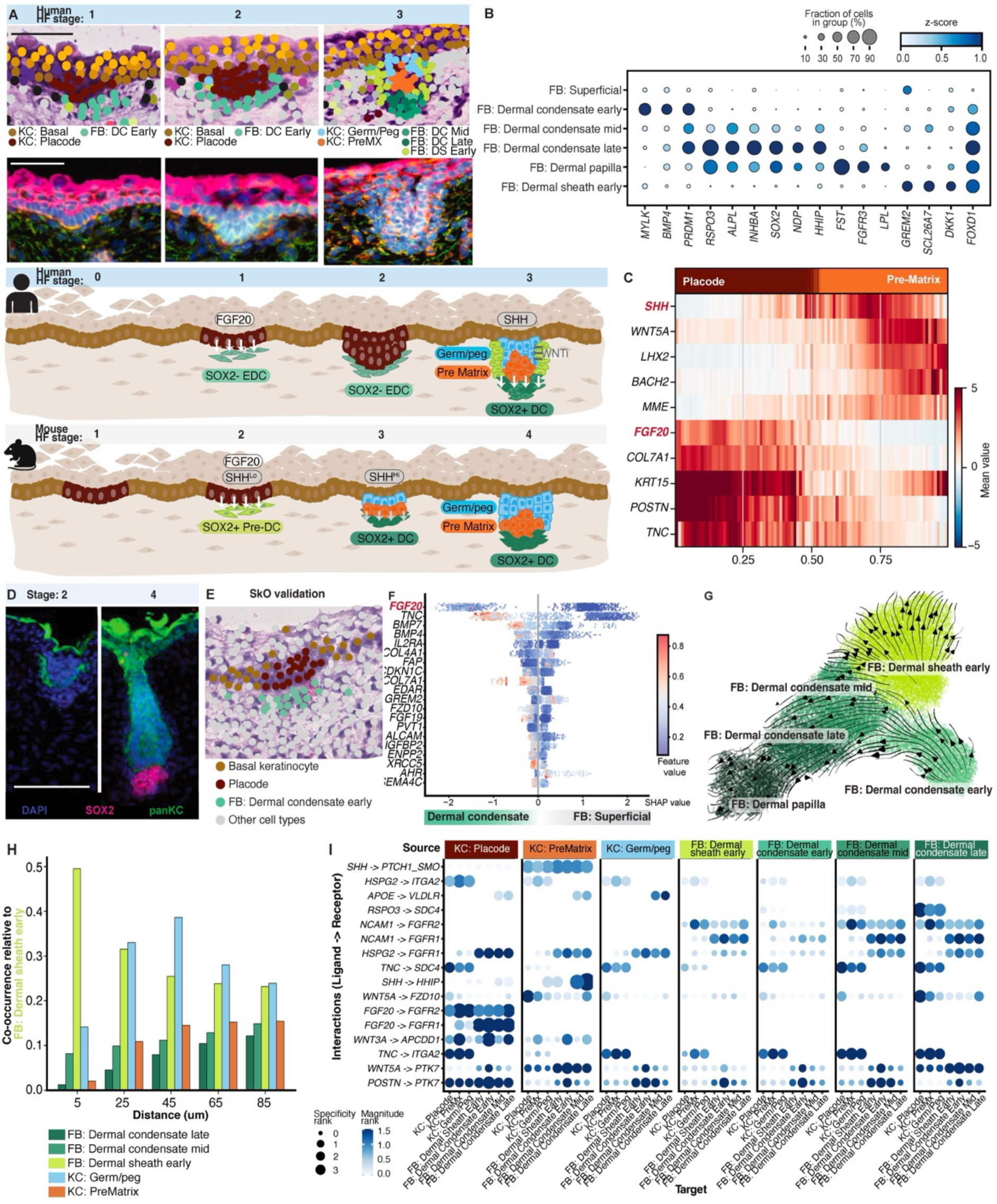
The Dermal Sheath and Dermal Papilla Both Descend From the Early Dermal Condensate. A Top: Placodes from Stages 2-4, top row: overlay of cell types with H&E, second row: Xenium boundary stains, third row: summary of new model for human HF initiation, bottom row: summary of murine HF initiation based on literature. Placode keratinocytes in Stages 1 and 2 are surrounded by dermal condensate Early cells, while in Stage 3 (right column) the placode keratinocytes have divided into PreMX and Germ/Peg keratinocytes. These PreMX and Germ/Peg keratinocytes are surrounded by two new fibroblast populations called EDS at the shaft sides, as well as dermal condensate mid/late on the bottom close to the PreMX. Bottom: New model for human placode initiation: together with the epithelial thickening, there is an EDC forming. After the placode has thickened, it is dividing both in the keratinocyte that splits into SHH+ PreMX and SOX9+ KC: Germ/Peg and in the fibroblasts, which differentiate into dermal condensate Mid/Late and EDS B EDC (MYLK, BMP4), dermal condensate mid/late (RSPO3, PRDM1, ALPL, INHBA, SOX2, NDP, HHIP), and EDS (GREM2, SLC26A7, DKK1) show expression of markers distinct from each other and the surrounding superficial fibroblasts. A common differentiator between the superficial and follicular fibroblasts is FOXD1. C Top genes associated with the change in expression between KC: Placode and KC: PreMX, showing loss of FGF20, COL7A1, TNC, and POSTN, and acquisition of SHH, LHX2, BACH2, MME, and WNT5A upon transition. D Immunohistochemistry confirming the presence of SOX2 in dermal condensate mid/late but not in early ones E Spatial location of placode and dermal condensate cells in SkO D76 placode F Shapley values of XGBoost prediction factors for environmental factors that drive EDC vs superficial fibroblasts (model roc_auc: 0.98) (every dot is a cell); color displays gene expression of the respective gene (feature). G Trajectory analysis of early follicular fibroblasts. The earliest progenitor, EDC (FB: Dermal Condensate Early), splits into two arms; one develops via the dermal condensate mid and late to the dermal papilla, and the other one to the EDS H Co-occurrence of early HF populations with EDS (FB: Dermal Sheath Early) 5-85 um I I Ligand-receptor based cell-cell communication analysis between early HF populations

As the species-specific delay between placode induction and *SHH*+ (KC: PreMX) cell fate acquisition could impact the timing of signaling molecules driving dermal condensate development, which is located below the PreMX, we next investigated the genes present during the transcriptional switch from placode to KC: PreMX. Genes associated with this developmental pseudotime were derived using a decoupleR univariate linear model (Methods), and included *SHH*, *WNT5A*, *LHX2,* and *BACH2* as the most upregulated and *FGF20*, *COL7A1*, and *TNC* amongst the most downregulated genes (Fig. 2C). The time interval between *FGF20* expression and the observed expression of signaling genes like *WNT5A* and *SHH* in the human placode is of importance, as these molecules induce dermal condensate in mice (*39*, *62*). The interval between FGF20 and SHH signaling in humans, together with the occurrence of *SHH*+ PreMX cells only after invagination of the placode, is distinct from murine KC: PreMX and dermal condensate formation, which is driven by simultaneous expression of *Fgf20 and Shh* (*63*).

To investigate the impact of the time interval between FGF20 and SHH, we investigated the dermal condensate identity before onset of SHH signaling. We observed molecularly distinct fibroblast condensates, expressing known murine dermal condensate genes such as *BMP4* (*32*), *PRDM1* (*64*), *FOXD1* (*63*), and genes not previously associated with the dermal condensate, such as *MYLK* (Fig. 2B), in proximity to the earliest detectable human placodes. We hereafter refer to these fibroblasts as Early Dermal Condensate (EDC) (Fig. 2A) as they exhibit the common dermal condensate morphology of elongated condensed fibroblasts but lack the expression of some previously reported murine dermal condensate genes, such as *SOX2* (*65*) and *HHIP* (*39*) (Fig. S6b). As *SOX2* is expressed in response to *SHH* signaling from the placode in mice (*66*), we investigated which genes might regulate SOX2 expression by assessing accessible chromatin around the *SOX2* locus in EDCs relative to adjacent superficial fibroblasts (Fig. S6c). This showed fewer open regions around the transcription start site of SOX2 in the EDC, but many more open regions up to 150kb upstream of SOX2 that are specifically open in the later stage of dermal condensate and might act as putative enhancer sites (Table S3). It also showed two regions specifically open in dermal sheath cells (Fig. S6c), and given that dermal sheath cells do not express SOX2, we speculate that they are repressor sites. To find TFs binding to these sites, we performed a CellOracle-based enrichment of TF binding sites in these peaks and found NR3C1, SP5, EGR1, and NRF1 to be most likely binding partners in these regions, suggesting that they regulate SOX2 expression during dermal condensate maturation (Table S3).

We validated the identity of the EDCs at the protein level by immunohistochemical staining, showing the morphology of the dermal condensate in Stage 2, but a lack of *SOX2* expression (Fig. 2C). Notably, *SOX2* expression was only observed once the follicles elongated (Stage 3 onwards), consistent with a recent study (*48*) (Fig. 2D). We additionally investigated if the human EDC stage was recapitulated in the SkO model and observed a *MYLK, FOXD1,* and *BMP4*-expressing mesenchyme population that is SOX2-negative, corresponding to prenatal scalp skin EDCs (Fig. 1D, 2E, S6d)(*46*). The presence of EDCs lacking expression of *SHH* response genes together with the time interval between *FGF20* and *SHH* expression in the placode are unique aspects of early human HF initiation events.

We next investigated the environmental signaling factors that induce the EDC population. Using the cell type-independent MISTy (*67*) framework employing tree-based predictions (XGBoost) from spatial views (see Methods), we identified *FGF20* expression in the placode to be the most predictive factor of human EDC fate acquisition (Fig. 2F), in keeping with reports from studies in mice (*34*, *68*). Further predicted factors were *BMP7* and *BMP4*, expressed by EDCs themselves, indicating that mesenchyme patterning independent of epithelial signaling might play a role in human dermal condensate formation, similar to that observed in mouse explants (*69*). Collectively, our findings demonstrate that human HF initiation is characterized by the concurrent formation of placode and dermal condensate, each occurring as a two-step process (Fig. 2A).

### The Dermal Sheath Arises From the Early Dermal Condensate

At Stage 3 of human hair development, we observed 2 distinct populations of hair-associated fibroblasts, one located deep at the base of the downgrowing HF, and one surrounding the HF shaft (Fig. 2A, right). The population at the base of the HF expressed canonical dermal condensate markers (Fig. 2B). The population surrounding the shaft expressed known markers of the dermal sheath, such as *SLC27A6* (*70*), but lacked the smooth muscle phenotype (*71*) (Fig. S6e) and was labelled as early dermal sheath (EDS). As the EDC, dermal condensate, and EDS co-locate in a cup-like structure around the downgrowing hair peg (Fig. 2B) and are transcriptionally distinct from the skin papillary fibroblasts (Fig. 2B), we hypothesized that EDC differentiated into dermal condensate and EDS, similar to murine dermal sheath cells that descend from dermal condensate (*72*). To investigate this, we performed trajectory analysis using the EDC cells as the starting point and observed, consistent with our hypothesis, a bifurcation that differentiates into two arms, one into dermal condensate continuing to the Dermal Papilla (DP), and the other into EDS (Fig 2G).

We next aimed to understand if the bifurcation in the fibroblast trajectory from EDC into dermal sheath and dermal condensate is coupled to the parallel bifurcation in the keratinocyte trajectory from placode into KC: Germ Peg and KC: PreMX. We thus quantified the co-location of KC: Germ/Peg and KC: PreMX with the fibroblast populations (dermal condensate mid/late and EDS) (Methods), which confirmed the visible co-occurrence of dermal sheath with KC: Germ/Peg and differentiated dermal condensate mid/late with KC: PreMX, suggesting their development is co-regulated (Fig. 2H, S6f).

To further investigate the interactions driving follicular keratinocyte and fibroblast differentiation during HF initiation, we performed ligand-receptor-based cell-cell communication analysis for these cells using LIANA+ (*73*) (see Methods). We observed co-expression of the hedgehog-pathway agonist *SHH* in the PreMX and *HHIP*, a hedgehog-pathway feedback inhibitor, in the dermal condensate mid/late (Fig. 2I). Corresponding genes of the hedgehog signaling pathway were upregulated in PreMX (*SHH*) and dermal condensate mid/late cells (*HHIP, SMO, PTCH1*) (Fig. S6g). These observations support the hypothesis that induction of the dermal condensate mid/late by KC: PreMX occurs through dynamic epithelial-mesenchymal cellular interactions, such as *SHH* signaling.

To investigate the timing of these cell-cell interactions and quantify their contribution to the EDC to EDS and dermal condensate mid/late differentiation, we first used the spatial location and morphology of cells in HFs to establish a temporal ordering, similar to a recent approach in mice (*74*) (Fig. S7a,b). Then, training a predictive model of lineage acquisition with a sliding window over the temporal ordering of hair development (Fig. S7a, Methods), we demonstrate that this lineage acquisition occurs between Stage 2 to Stage 3 (window with the center of 30) (Fig. S7c) and that the most important predictive features for dermal sheath over dermal condensate lineage acquisition include several Wnt inhibitors such as *DKK1,* and *GREM2* (Fig. S7d). In mice, DKK1 was reported in similar locations (*30*, *62*), but also in the dermal condensate (*75*), supporting the hypothesis of its involvement in dermal sheath formation. Together, the expression of these WNT factors in the EDS with their implication in a predictive model of EDS and dermal condensate specification suggests that Wnt inhibition is an important factor in dermal sheath specification (Fig. 2A).

### Dermal White Adipose Tissue Arises Surrounding Stage 10 Hair Follicles

In mice, a thin sheet of striated muscle called the panniculus carnosus acts as the classical endpoint of hair growth, but humans largely lack this structure (*76*). We therefore aimed to understand the cues that govern HF downgrowth into the lower dermis, which is known to extend deeper after birth (*77*). Developing scalp skin is stratified into 4 transcriptionally distinct layers of fibroblasts that correspond to the papillary dermis (superficial), upper or lower reticular dermis, and hypodermis (Fig. S8a,b). Additionally, we identified genes expressed in gradients along the vertical axis of the section (Fig. S8c, Table S4, Methods). Stage 10 HFs were observed at the intersection between the lower reticular dermis and the hypodermis, which is more superficial than in mice (Fig. S8def).

The lower reticular dermis and hypodermis consist of fibroblasts, blood and lymphatic vessels, and adipose tissue (*78*). Dermal white adipose tissue is known to surround adult HF bulbs (*79*). In mice, the origin of adipocytes was traced to reticular and hypodermal fibroblasts (*80*), while in humans the origin of different adipocyte populations has not been shown. Starting from 14-16 PCW, we identified FB: PreAdipocyte cells (Fig. S8g) expressing adipocyte marker genes such as *ADIPOQ*, *LPL,* and *CIDEC* in the lower reticular dermis (Fig. S8hi). Additionally, using footprint-based enrichment analysis (Methods), we detected activity of classic adipocyte TFs such as Liver X Receptor (*NR1H2*, *NR1H3*) and *PPARG* (Fig. S8j), and Adipogenesis as the most prominent gene programme in FB: PreAdipocytes (Fig. S8k). Starting from 17 PCW, the uniform distribution of FB: PreAdipocyte cells in the lower reticular dermis transforms into a majority of cells locating around downgrowing hair bulbs (Fig. S8ef). Cell-cell communication analysis using LIANA+ identified several putatively inhibitory interactions between the dermal papilla stalk, dermal sheath, and the adipocytes around the hair bulb, including *TGFB2-TGFBR3* and *SLIT2-ROBO1,* suggesting adipocytes might play a role in regulating the depth of hair growth (Fig. S8l, Methods). We further observed that the HFs grow through the dermis regions in the SkO where the adipocytes are absent, supporting the potential inhibitory role of adipocytes in hair growth (Fig. S8m).

### Medio-lateral Differentiation of Epithelial-derived Hair Follicle Layers

In addition to its vertical elongation, the HF differentiates medio-laterally during morphogenesis. The outermost epithelial layer is the ORS, which descends from the KC: Germ Peg (Fig. 1D). The next layer inward is the companion layer (CL), expressing *KRT79* (*81–83*), a cornified layer of epithelium building the hair canal that separates the ORS from the innermost structures of the Inner Root Sheath (IRS) and the hair shaft(*81–83*).

CL and IRS have been previously identified in humans (*47*) and mice (*57*, *74*, *82*); however, it is unclear at which HF stage they form in humans. Between Stages 4 and 10, we observed 3 transcriptionally distinct populations of cells radiating outwards from the hair shaft at the centre of the HF to the ORS (S9a). We annotated these three populations as IRS, CL, and PreCL based on spatial location and marker gene expression. The IRS and CL are found at Stage 7 onwards, while the PreCL is observed earlier at Stage 4 (Fig. 3A). The IRS is observed above the KC: Matrix (IRS progenitor) in Stage 7 of HF development and expresses well-known IRS markers such as *KRT71*, *KRT27* (Fig. 3A, B, C). Up to 19 PCW, IRS cells occupy only the lower third of the elongated follicle, extending from the hair bulb to the point where the hair shaft emerges, whereas in mice they extend more superficially, reaching well above the hair bulge and sebaceous gland (*36*, *39*, *84*) (Fig. 3A, B). The SkO HF also has an IRS with a similar location to prenatal scalp HF (Fig. S9a).

**Fig. 3:**
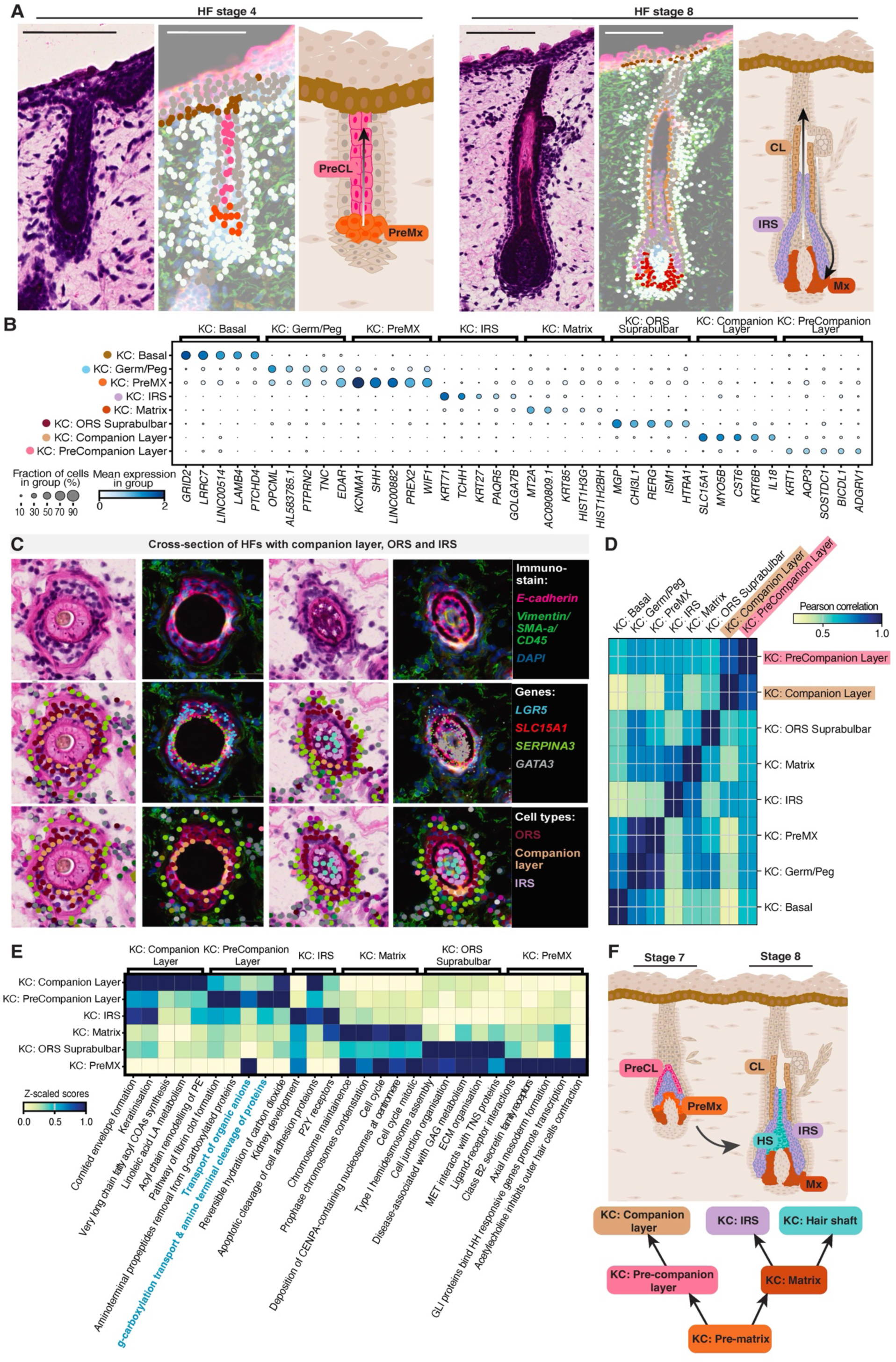
The Companion Layer descends from early suprabasal cells. A Annotation of Stage 4 and 8 HFs depicting the spatial location of KC: Matrix (brown), IRS (purple), CL (beige) cells, and their progenitors. B scNuq-RNA markers of cell types presented in A and B. PreMX cells strongly express SHH, which is lost in the more mature KC: Matrix. The KC: Matrix itself expresses mostly proliferation markers. The CL expresses KRT6B as well as SLC15A1 and CST6, compared to the PreCL that specifically expresses SOSTDC1 and KRT1. The IRS expresses well-known markers such as KRT71 and KRT27. C Top row: Cross-sections of HF that show the expression of SLC15A1 (red) as well as SERPINA3 (green) as markers of the CL, as well as GATA3 (grey) as IRS, and LGR5 (blue) as ORS marker. Middle row: Cross-section alone, between the IRS and the CL, a clear gap is visible. Bottom row: cell type annotation. D ScNuc-RNA correlation KC: Matrix, showing higher correlation (r = 0.81) between PreCL and CL than with any other cell type in the proximity. E Reactome pathway analysis showing enhanced cornification in the CL and similarity between PreMX and PreCL. F Family tree of PreMX and KC: Matrix descendant cell types. PreCL and CL separate early as descendants from PreMX, while the IRS and Shaft originate from the KC: Matrix.

The CL is physically separated from the IRS in the medio-lateral axis and expresses marker genes of the CL, such as *KRT6A/B* and *CST6* (*81*) (Fig. 3C, S9b). Furthermore, it lines the inside of the ORS (Fig. 3B) (*81*) as reported in murine lineage tracing experiments (*57*, *85*) and in adult human HFs (*82*, *83*). Corresponding CL cells were found in our human adult and SkO HF datasets (Fig. S9c,d). Notably, the CL appeared at the same time as the KC: Matrix was formed, and as such cannot be KC: Matrix-derived (Fig. 3A, D, S9a). Taken together, this shows that the developing IRS (KC: Matrix-derived) and CL (not KC: Matrix-derived), arising at Stage 7 and 8 of HF development, have distinct cellular origins, as shown in mice (*57*).

The PreCL lines the inside of the KC: Germ/Peg, similar to CL progenitors in mice (*57*) (Fig. 3A), and was transcriptionally similar to the CL and had similar open chromatin correlation (Fig. 3E, S9e,f). In the cropped HFs (Table S2), it was consistently observed in HFs in which the CL has not yet formed (Fig. 3A). In mice, the PreCL originates from PreMX (*57*, *85*). Consistent with this, our ATAC data showed that the PreCL was more correlated with KC: PreMX (0.35) than KC: Germ/Peg (0.19). Characterisation of the PreCL transcriptome revealed it was enriched with suprabasal keratinocyte markers, expressing *KRT1* (Fig. S10a,b) and *CDKN2B* (Fig. S10a). Gene set enrichment analysis using Reactome gene sets (Methods) highlighted functions associated with transport, including anions, which are needed for lumen formation (*86*, *87*), aligning with a potential role in hair canal formation. Consistently, PreCL specifically expressed *FOXN1* and *SOSTDC1* (Fig. S10b), which have been associated with hair canal defects (*88*) and regulation of HF size (*89*) in murine studies. This expression pattern was also observed in the SkO, suggesting shared programs of hair canal formation between the two systems (Fig. S10c). Taken together, this shows that the terminal keratinocyte differentiation programme is shared between PreCL and suprabasal interfollicular keratinocytes and suggests the PreCL, likely derived from KC: PreMx, to be involved in hair canal formation.

### Inner Root Sheath Lineage Acquisition Occurs in the Distal Tip of the Bulb

The main function of the KC: Matrix is to produce the hair fiber, which consists of enucleated hair shaft cells that ascend vertically in the center of the follicular unit, surrounded by the IRS (Fig. 4A). This upward movement (y-axis position) of the IRS and hair shaft cells is marked by expression of several genes such as the iron exporter Ferroportin (*SLC40A1*) and Heme Oxygenase 1 (*HMOX1*), as well as water transport (*AQP3*) and cell-cell adhesion (*LRRC15*) (*90*) (Fig. S11a,b). We also detected activity of transcription factors such as KCNIP3, IRF5, and TBPL2 (Fig. S11c-d, Methods). We used the anatomical position of the IRS and hair shaft to identify marker genes that are distinct for the hair shaft (*KRT85*, *HOXC13*, and *CTSC*) (Fig. 4B, S11a, Table S2). Using this annotation, we identified that the first hair shaft forms at Stage 8 following IRS formation at Stage 7 (Fig. 1D).

**Fig. 4:**
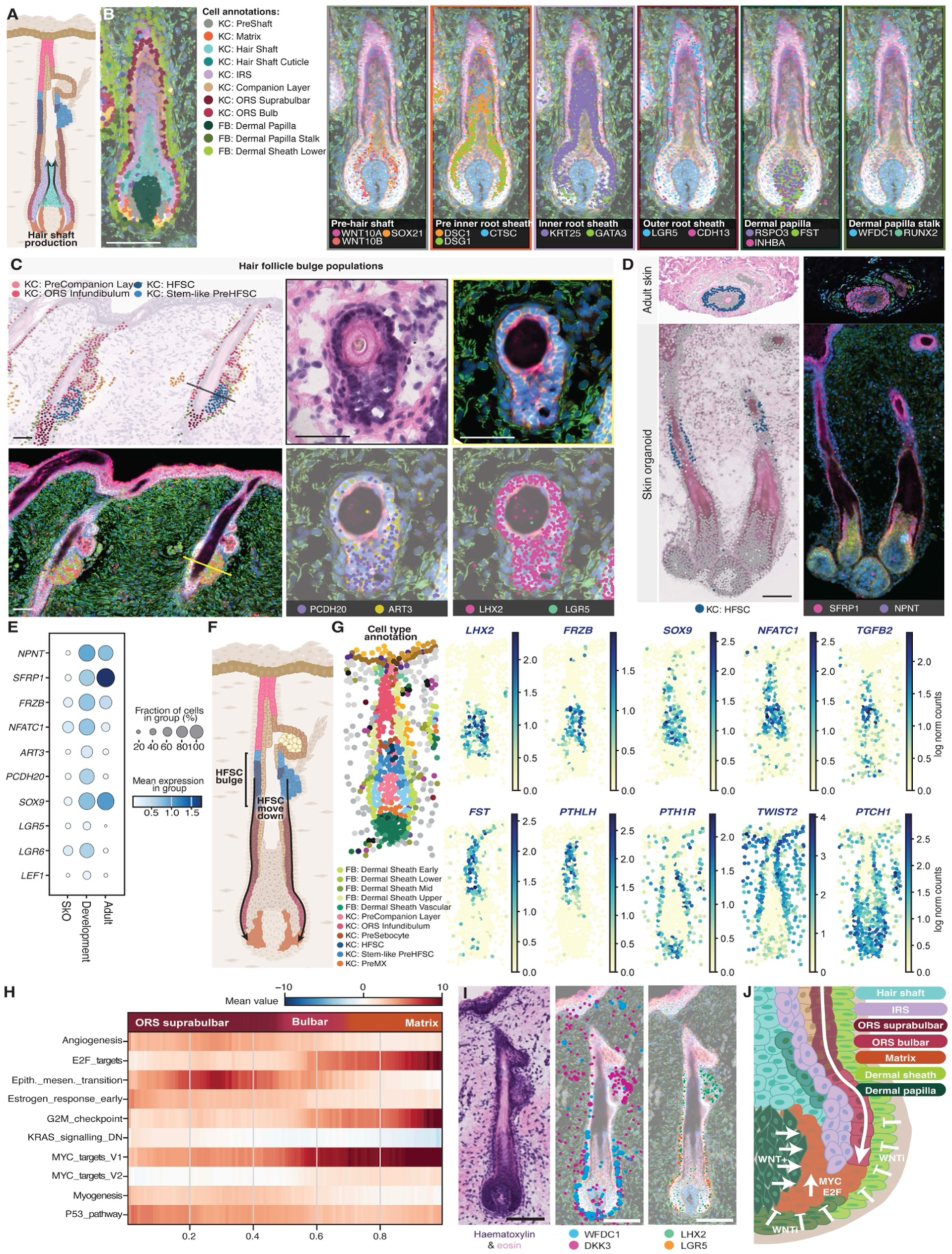
Lower ORS cells transition from dormant to activated in the dermal sheath niche. A Cartoon highlighting the main cell types involved in hair fiber production. B Markers of bulb populations compiled from analysis of spatial and snRNA-seq data. C Left: annotated cell types overlaid with the H&E, bottom middle boundary stains with transcript location of PCDH20 and ART3. Right: cross-section of the bulge; outer bulge cells seem to have apical-basal polarization as well as expression of ART3 and PCDH20 (top right) and LHX2 and LGR5 (bottom right) D Top: Spatial location of adult HFSC cells. Bottom: Location of label-transferred HFSC in the SkO E HFSC Marker expression of human developing scalp hair, adult hair, and SkO hair F Cartoon showing the populations involved in the downward movement and KC: Matrix replenishment G Top genes associated with skin depth (y-axis) in ORS keratinocytes (LHX2, FRZB, SOX9, NFATC1, TGFB2, FST, PTHLH) and dermal sheath fibroblasts (PTH1R, TWIST2, PTCH1) highlight two transcriptional segments where, at the border, preHFSCs and PreSebocytes are detectable (right) H Hallmark enrichment along the axis from ORS suprabulbar to Bulbar to KC: Matrix, constructed by computing a pseudotime starting at the ORS suprabulbar. Upon transition from ORS bulb to KC: Matrix, we can see a strong increase in G2M checkpoint, E2F, and MYC targets, all related to a strong increase in proliferation. I Location of stem cell markers (LGR5, LHX2) in ORS and Wnt inhibition (DKK3, WFCDC1) in lower dermal sheath J Cartoon showing the location of the area where the transition from ORS to KC: Matrix is facilitated by a switch from WNTi inhibition to WNT activation

To identify the spatial location of the lineage segregation between the IRS and hair shaft, we assessed transcriptional heterogeneity of the bulb epithelial cells and observed three populations that differentially express marker genes of the ORS, IRS, and the hair shaft (Fig. 4B). As they represent the likely descendants of ORS and progenitors of IRS and hair shaft, we annotated them as postORS (*LGR5, THBS1,* and *TNC*), preIRS (*DSC1, DSG1, NOTCH3*), and preShaft (*SOX21*) based on their gene expression and spatial location (Fig. 4B, S12a). The transition point from postORS to preIRS and preShaft is located at the distal tip of the bulb at the border between the dermal papilla and dermal papilla stalk, suggesting that lineage segregation occurs here (Fig. 4B).

To investigate the regulators of the fate decision between IRS and hair shaft, we applied our MISTy approach (Fig. S12b). We found that expression of *FREM2, COL17A1,* and *PRXL2A* around PreShaft cells predicted hair shaft fate, while *NOTCH2* and *NOG* were the most predictive factors for the IRS fate, and this prediction was supported by the spatial location of the respective genes (Fig. S12c-d). Supporting the role for NOTCH, we observed NOTCH Hallmark and Reactome pathway enrichment in PreIRS-specific cells (Fig. S12ef), and upregulation of NOTCH 1/2/3 and NOTCH2LA expression (Fig. S12f). In the SkO, *NOTCH2* is expressed in the IRS along with the classical IRS markers, suggesting that NOTCH2 has a functional role in lineage segregation due to its conserved expression between systems (Fig. S12g). Together, this shows that IRS lineage acquisition is characterized by NOTCH expression and activity.

### Developing Scalp Bulge Cells Are Morphologically and Transcriptionally Distinct From Outer Root Sheath

Hair fiber production comes with design requirements that are reflected in the geometry of the HF to allow controlled growth (hair cycling) but withstand dysregulation to prevent hyperplasia or potentially accumulation of somatic mutations. This is achieved by producing a stem cell reservoir in a structure called the bulge (*7*, *8*, *91*), which allows the HF to go through a cycle where the entire lower region of the HF is renewed (*23*). In developing human scalp HFs, the bulge develops together with the ORS from the KC: Germ/Peg and first appears positioned at the vertical center of the ORS in Stage 7 HFs (Fig. 1D).

While the developing human scalp bulge is histomorphologically distinct (Fig. 4C), the molecular identity and drivers of its development remain unknown. Differential gene expression analysis of bulge cells identified HFSCs that were characterized by the expression of ADP-ribosyltransferase 3 (*ART3*), Nephronectin (*NPNT*), Secreted Frizzled-Related Protein 1 (*SFRP1*), and Protocadherin 20 (*PCDH20*) (Fig. 4c, S12h). *Npnt* and *Sfrp1* are associated with arrector pili muscle induction (*24*) and Wnt inhibition (*92*, *93*) in the murine bulge. *ART3* and *PCDH20,* two genes with roles in spermatogenesis and cell-cell adhesion, are unique to human developmental HFSCs and restricted to the bulge; therefore, we investigated the chromatin state of *ART3* and *PCDH20* to understand how these genes are regulated. Surprisingly, there was no open chromatin at the canonical transcription start site of *ART3* in any cell type (Fig. S12i). However, we could confirm a bulge-specific peak at the alternative transcription start site of the isoform ART–226, suggesting that this alternative isoform is likely expressed in HF. *PCDH20* exhibited several open regions 220 kb downstream and co-accessible with the *PCDH20* transcription start site in *PCDH20-expressing* cell types, suggesting they are enhancers of *PCDH20* expression (Fig. S12j). Collectively, this suggests *ART3, NPNT, SFRP1*, and *PCDH20* as transcriptional markers of the developing human scalp bulge. Future studies are required to define the functional roles of these proteins in human bulge formation.

Although adult HFs have a visible bulge structure housing HFSCs, the SkO did not show a histomorphologically distinguishable bulge (Fig. 1D). However, HFSCs were observed in the ORS at roughly two-thirds of the HF down from the epidermis, where a bulge would be expected to form (Fig. 4D, Methods). Although the SkO HFSCs expressed the canonical markers *NFATC1* (*44*) and *LGR6* (*40*) of bulge cells, they lacked classical HFSC markers such as *SFRP1* and *NPNT*, suggesting immaturity and aberrant stem cell development in the SkO compared to *in vivo* (Fig. 4D, E).

### Lower ORS Harbors Dormant Stem/Progenitor Cells

In addition to enabling the hair cycle, the bulge cells migrate down the ORS (*94*, *95*) and replenish the transit amplifying cells (*96*), similar to a conveyor belt (Fig. 4F). This downward cell movement has been appreciated in the adult anagen HF (*94*, *95*). However, whether stem cell identity is maintained at the level of the lower ORS (KC: ORS Bulbar and KC: ORS Suprabulbar) has not been confirmed in development. We find that despite appearing histomorphologically uniform, ORS cells are split into two segments by transcription and position (Fig. 4G, Table S5, Methods). The upper segment starting at the middle of the ORS and extending up to the epidermis (termed ORS Infundibulum) showed high expression of Wnt/Bmp modulator genes Follistatin (*FST*) and Parathyroid Hormone Like Hormone (*PTHLH*) (Fig. 4G). In contrast, the lower segment containing the suprabulbar and bulbar ORS exhibits expression of classical HFSC genes such as *LHX2* (*43*) and *LGR5* (*41*), and a similar fraction of cycling cells to the bulge HFSCs (Fig. S13c). These segments are also observed in SkO Stage 5 HFs (Fig. 1D, S13b). Further stem cell markers such as *SOX9* (*9*) and *NFATC1* (*44*) were present in the lower ORS segment and central region of the ORS (Fig. 4G, S13a). Together, these findings suggest that the lower ORS, including the suprabulbar and bulbar ORS, harbours HFSCs that transition from the Bulge to the KC: Matrix.

We next asked how the lower ORS cells transition from dormant stem cells towards proliferative KC: Matrix cells. We calculated a pseudotime starting from the suprabulbar ORS to the bulbar ORS and the KC: Matrix (Fig. S13d, Methods) and identified a positive association of the hallmark G2M checkpoint with the pseudotime, as well as the upregulation of *E2F* and *MYC* target genes (Fig. 4H). The upregulation of *E2F* transcription factors was further supported by a footprint-based transcription factor (TF) activity estimation (Fig. S13e, Methods). Our findings suggest that the HFSCs in the ORS acquire proliferative capacity through *E2F* and *MYC* target gene expression in a similar mechanism to how interfollicular epidermal stem cells become transit amplifying cells(*97–99*). In epidermal stem cells (*100*) and other cell types (*101*, *102*), *MYC* is regulated by several upstream pathways, including MAP/ERK, Wnt, and JAK-STAT(*103*). To understand its upstream regulation in the HF KC: Matrix, we performed pathway enrichment along the ORS-to-KC: Matrix trajectory and observed Wnt-betaCatenin signaling to be consistently upregulated in the KC: Matrix (Fig. S13f), suggesting Wnt-betaCatenin as the *MYC* activator in the KC: Matrix.

Stem cells reside in niches (*104*) that keep them dormant. Accordingly, during the transition from HFSC to proliferative KC: Matrix cells, ORS stem cells need to be kept in a stem cell state before Wnt signaling exposure for activation. The only cell type in the vicinity of the lower ORS that could provide such a niche is the mesenchymal dermal sheath, which was speculated to be involved in stem cell maintenance in mice (*105*). We thus investigated stem cell niche regulators related to Wnt signaling along the “conveyor belt”. In addition to the well-known Wnt activating signals such as *WNT5A*, *CTNNB1*, and *RSPO3* originating from the dermal papilla, we could detect several Wnt inhibitors (*WIF1, DKK1/3, SFRP2)* and modulators (*WFDC1*, *LGR6*), expressed in the dermal papilla stalk and lower dermal sheath, which surrounds the stem cells in the lower bulbar ORS (Fig. 4I, S13g). The SkO lower dermal sheath and dermal papilla stalk also showed expression of *DKK3* and *WFDC1* but lower levels of *SFRP2,* suggesting a similar mechanism of Wnt inhibition in SkOs (Fig. S12h). Wnt inhibition by the lower dermal sheath along the downward travel of the lower ORS cells prior to Wnt activation by the dermal papilla is a plausible mechanism for how the stem cells keep their dormant state before entering the KC: Matrix stage.

To dissect further genes involved in the transition from HFSC-ORS to KC: Matrix, we revisited the KC: Matrix subclusters (Fig. 4B). Notably, the postORS expresses *LGR5* (Fig. S12a), which can potentially bind to *RSPO3* expressed on dermal papilla cells (Fig. 4B) to increase WNT sensitivity (*106*). Our findings suggest a switch from Wnt inhibition to activation upon exposure to the dermal papilla that upregulates *MYC*/*E2F* expression and subsequent acquisition of proliferation capacity (Fig. 4J). This is in line with the observation that depletion of Wnt (*107*) or beta-catenin (*108*) in the dermal papilla leads to reduced KC: Matrix proliferation in mice and altered KC: Matrix formation, where Rspo3 and Lgr5 have been upregulated or depleted (*109*, *110*). However, unlike in mice, where expression gradients in the dermal papilla are reported to account for IRS specification (*111*), in the developing human dermal papilla Wnt activators were homogeneous and constitutively expressed (Fig. 3B). Taken together, this suggests a new function of the lower dermal sheath as a stem cell niche regulator that restrains migrating HFSCs in a dormant state through Wnt inhibition before HFSCs are activated into transit-amplifying cells in the KC: Matrix.

### Arrector Pili Maturation is Driven by Cardiomyocyte-like Myocardin-Serum Response Factor Transcriptional Activation

Closely located to the Bulge is the attachment site for the arrector pili muscle, a smooth muscle associated with piloerection (also known as “goosebumps”)(*112*), thermoregulation (*113*), and hair loss (*114*)(Fig. 6A). While the attachment and induction processes on the bulge side are well studied in mice(*24*), and Merkel cells are thought to facilitate the attachment of the arrector pili muscle on the epidermal site in humans (*115*, *116*), little is known about the maturation of the arrector pili muscle itself. In human prenatal scalp skin, we observed arrector pili muscle cells attached to the HF in Stage 6 HFs at PCW 14 (Fig. S14ab) expressing Actin Alpha 2 (*ACTA2*, α-SMA), Transgelin (*TAGLN*), Integrin alpha-8 (*ITGA8*)(*24*), and the canonical receptor alpha-1A adrenergic receptor (*ADRA1A*)(*117*, *118*) (Fig. 5AB, Fig. S14c). Immunohistochemical staining confirmed ACTA2 expression in the arrector pili muscle (Fig. S14d).

**Fig. 5:**
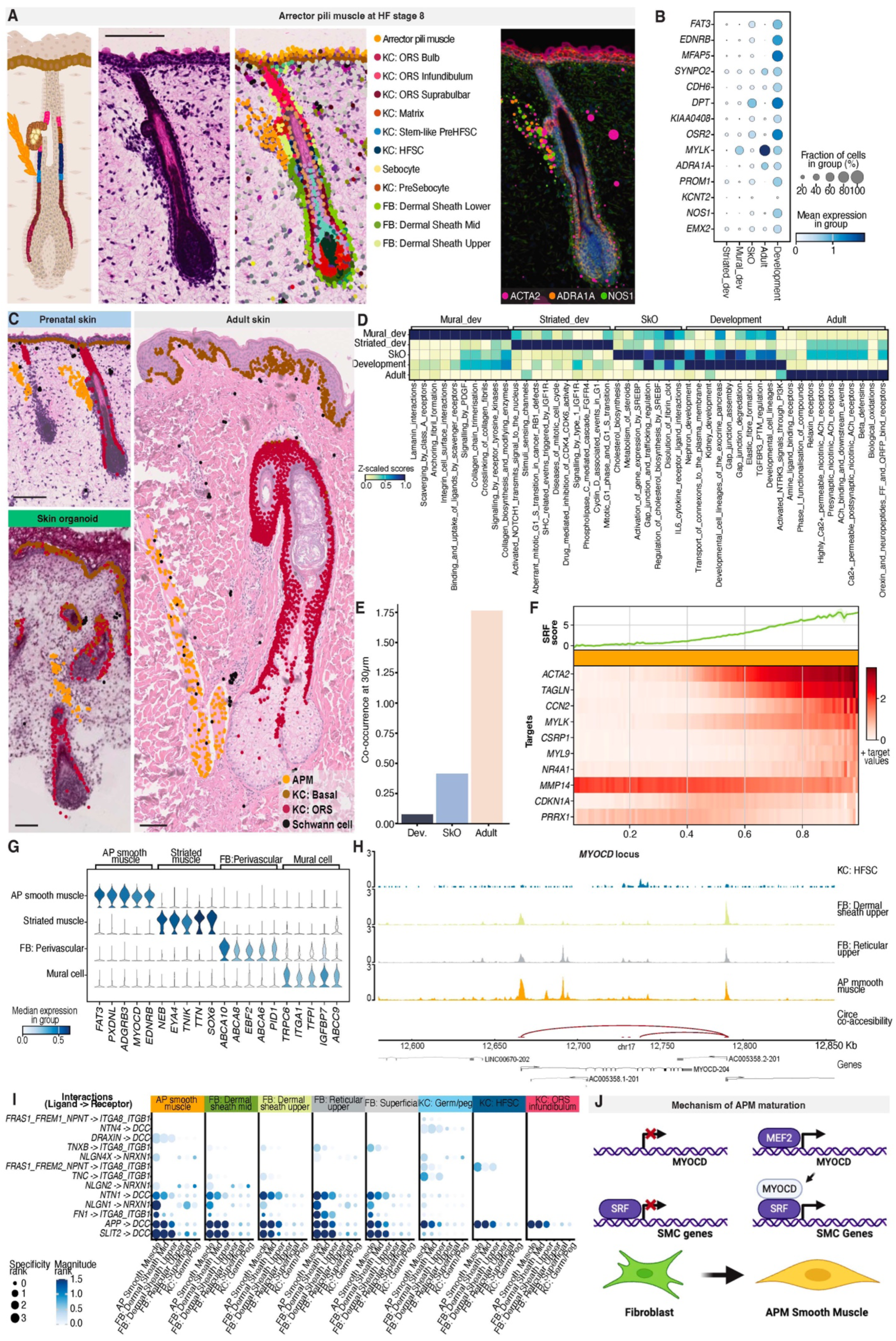
Arrector pili muscle maturation exhibits Myocardin-enhanced Serum Response Factor activity. A Left: Cartoon highlighting the location of the arrector pili muscle and different ORS populations, Middle: Spatial location of arrector pili muscle cells shows proximity to the bulge area below the sebaceous gland, Right: Spatial location of arrector pili muscle Marker transcripts ACTA2, ADRA1A, and NOS1. B Dotplot of top markers of developmental arrector pili muscle across adult and SkO, as well as other developmental muscles. C Adult arrector pili muscle shows colocation with Schwann cells, indicating innervation that is not observed in arrector pili muscle of developmental or SkOs. D Reactome pathway enrichment showing enrichment of neuronal signaling pathways in the adult arrector pili muscle. E Co-occurrence of arrector pili muscle and Schwann cells at 30 um in arrector pili muscle from adult, developmental, and SkO. F SRF and downstream target activity along the arrector pili muscle maturation trajectory constructed by pseudotime inference (correlation to real time in PCW: r=0.52). G ScNuc-RNA gene expression markers of arrector pili muscle and other muscles in development. H ATAC coverage around the myocardin locus showing specific open regions around the myocardin transcription start site. I LR-based cell-cell communication analysis between arrector pili muscle cells, related FBs, and HFSCs. J Schematic of arrector pili muscle Maturation: In normal fibroblasts, neither MYOCD nor smooth muscle genes are expressed. During arrector pili muscle maturation, MEF2A/C activates myocardin (MYOCD), which binds to the serum response factor that, in turn, activates smooth muscle genes.

**Fig. 6:**
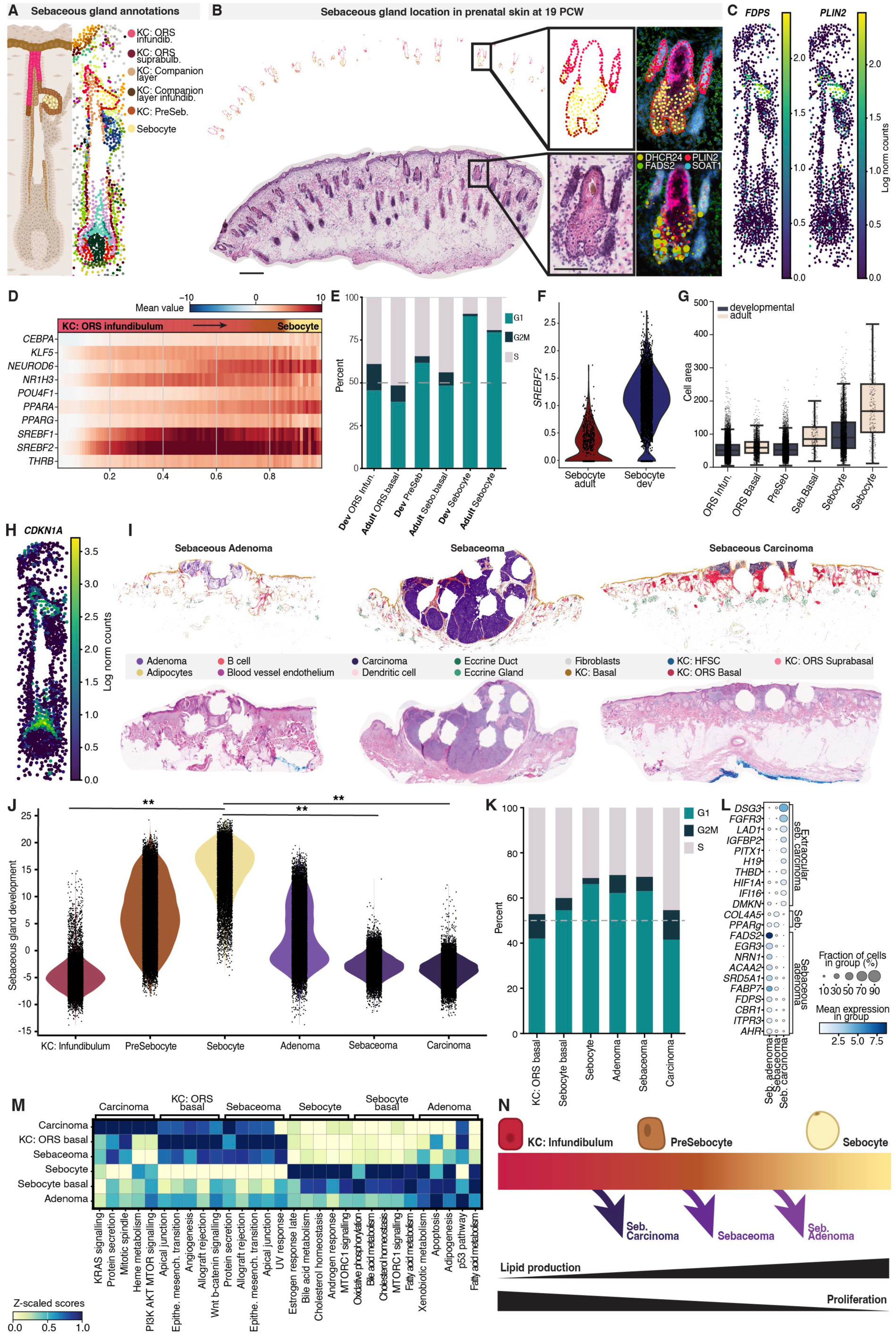
Sebaceous gland development is characterized by a shift to lipid production and cell cycle arrest, which is lost in sebaceous tumors. A Cartoon showing the spatial location of PreSebocytes and Sebocytes, B Tissue location of PCW19 sebaceous glands, their cell types, and markers such as DHCR24, PLIN2, FADS2, and SOAT1 C Expression of sebaceous gland markers such as FDPS and PLIN2 is specific throughout the whole HF section. D TF activity enrichment along the developmental trajectory from ORS infundibulum to PreSebocyte and sebaceous gland, constructed by pseudotime starting at ORS infundibulum. The strongest association with the pseudotime is exhibited by SREBF1/2 and PPARA. E Logarithmized counts per thousand SREBF2 expression between developmental and adult Sebocytes F Cell cycle phases between the ORS infundibulum, PreSebocytes, sebaceous glands, and their counterparts in the adult sebaceous gland. Proliferation (percent of cells in S or G2M phase) is generally higher in adult counterparts G Cell area between ORS infundibulum, PreSebocytes, sebaceous glands, and their counterparts in the adult sebaceous gland. Adult cells are consistently larger than their developmental counterparts H Expression of CDKN1A (P21) along the hair section. Strongest areas of expression are the sebaceous gland and hair shaft. I Location and H&E morphology of cell types across three types of sebaceous tumors ordered after severity. While the sebaceous adenoma consists of mostly normal-looking Sebocytes, the sebaceoma cells look aberrant. The sebaceous carcinoma phenotype is more commonly located on the surface of the epidermis and looks more KC-like J Enrichment of sebaceous gland developmental program along non-tumor sebaceous glands and their tumorigenic counterparts shows an increase in enrichment in more mature non-tumor phenotypes, while the tumor cells are less enriched with their severity and non-typicality. Significance was assessed by weighted least-squares regression (inverse-variance weighted). P values were adjusted using the Benjamini–Hochberg FDR procedure. *FDR < 0.05, **FDR < 0.01. K Cell cycle phases of sebaceous tumor show an increase in proliferation with tumor severity and sebaceous gland developmental program signature L Marker genes of tumor cells show expression of lipid production genes such as PPARG in the non-malignant cell states, while the carcinoma cells undergo hypoxia, as well as the expression of HIF1A M Hallmark enrichment of tumor and non-tumorigenic cell types shows activity of fatty acid and cholesterol pathways in the least severe adenoma, while the more severe sebaceoma and carcinoma have an increase in epithelial-mesenchymal transition. The carcinoma shows specific upregulation of cell cycle-related pathways, such as the G2/M checkpoint N Overview of the sebaceous gland developmental program that links fatty acid and cholesterol production to a halt in the cell cycle. The earlier a cell escapes this program, the more proliferative, severe, and less sebaceous gland-like the tumor will be.

Beyond these characteristic genes, developing arrector pili muscle cells also express Nitric Oxide Synthase (*NOS1*), an enzyme producing nitric oxide that is known to relax smooth muscle cells in the cardiovascular system (*119–122*) (Fig. S14c). The potential production of nitric oxide poses the question of whether the arrector pili muscle is non-contractile when the fetus is bathed in amniotic fluid *in utero*, and is only active after birth when thermoregulation becomes critical. We therefore compared the transcriptome of the developing arrector pili muscle with adult skin and SkO samples, which confirmed that *NOS1* expression is restricted to prenatal skin (Fig. 5B, S14e, Methods).

In addition to contractility, an important feature of a functional arrector pili muscle is its innervation (*123*). In adult skin arrector pili muscle, we observed Schwann cells intercalated with the arrector pili smooth muscle cells, indicating innervation of the arrector pili muscle (Fig. 5C, right). In contrast, prenatal skin and SkO arrector pili muscles had few intercalated Schwann cells (Fig. 5C, left, 6E). This difference is also reflected by a Reactome pathway activity inference, in which adult arrector pili muscle cells activate pathways related to nicotinic acetylcholine (nACh) receptors and amine receptors that are downregulated in prenatal and SkO arrector pili muscle, as well as other developmental muscle cells (Fig. 5D). Collectively, our findings suggest that even though the prenatal hair has reached maturity at 10 PCW, prenatal arrector pili muscles are non-contractile, as indicated by the expression of *NOS1* and the lack of innervation, consistent with the expectation of arrector pili muscles’ role in thermoregulation, a function being required only after birth.

Murine studies suggest that the arrector pili muscle progenitors arise from mesenchymal cells around the bulge that mature towards a smooth-muscle-like phenotype upon exposure to nephronectin (*NPNT*) and other signals from the bulge (*24*). Given that both the arrector pili muscle and dermal sheath have contractile functions, we investigated whether these cell populations have a shared origin. Our data demonstrates that dermal sheath fibroblasts and the arrector pili muscle cells have similar transcriptomes, supporting the suggestion of shared origin (Fig. S14f). As the SkO lacks an *NPNT*-expressing bulge, the arrector pili muscle anchor site is lower on the vertical axis than *in vivo* (Fig. 5C), confirming the phenotype from an Npnt-KO study in mice (*24*). We then wondered what drives the maturation once dermal sheath cells are committed to the arrector pili muscle fate. Using inferred trajectory and pseudotime analysis, we investigated the TFs that drive the maturation of fibroblasts into arrector pili muscle (Fig. S14g). The pseudotime was correlated with gestational time (Pearson r: 0.52)(Fig. S14h). Among the top pseudotime-associated transcription factors were TFs involved in heart development, *KLF13* (*124*), *HEY2* (*125*), TCF21 (*126*), *HHEX* (*127*), and, most notably, the serum response factor (*SRF*)(*128*) (Fig. 5F, S14i). *SRF* response genes include many smooth-muscle-associated genes, including *ACTA2* and *TAGLN*, that are upregulated in the arrector pili muscle (Fig. 5F).

*SRF* is a ubiquitously expressed transcription factor that works together with transcriptional co-activators such as myocardin to drive a contractile phenotype in smooth muscles (*129*) and cardiomyocytes (*128*). In concordance with the literature, we see low ubiquitous expression of *SRF* in all cell types (Fig. S14j) and specific expression of myocardin in arrector pili muscle cells, differentiating it from all other muscle types in the dataset, including striated and vascular smooth muscles (Fig. 6G). Additionally, using a footprint-based approach to assess open chromatin (see methods), we identified arrector pili muscle-specific activity of myocardin (MYOCD) in developmental muscle cells (Fig. S15a), which has also been reported in mice (*130*). Similarly, *MYOCD* is expressed in the SkO arrector pili muscle (Fig. S15b). To confirm the presence of myocardin in the arrector pili muscle cells, we stained sections of fetal skin at PCW 17 and a whole-mount of day 145 SkO with anti-myocardin antibodies and observed expression of myocardin in arrector pili muscle cells (Fig. S14d, S15c). Together, these analyses suggest that arrector pili muscle maturation is driven by myocardin and SRF activity.

To investigate the upstream induction mechanism of myocardin, we searched for co-accessible peaks within the myocardin transcription start site using Circe (*131*) (Methods). We could observe several regions only open in arrector pili muscle cells, including regions annotated as enhancers by ENCODE (*132*), suggesting that they might have a direct regulatory role in myocardin expression (Table S6)(Fig. 5H). To understand what transcription factors bind in these arrector pili muscle-specific open regions, we looked for enriched transcription factor motifs in those peaks with CellOracle (*133*) (see Methods). The top two transcription factors that could be associated with the co-accessible peaks were *MEF2C* and *MEF2A*, two transcription factors known to regulate myocardin in heart development (*134*, *135*) and murine arrector pili muscle (*130*) (Fig. 5H, Table S7). Together, the pseudotime, chromatin accessibility and co-accessibility analyses, combined with the known regulatory relationship between MEF2 and MYOCD in other muscle contexts, support a model in which MEF2 promotes MYOCD expression, enabling MYOCD together with the broadly expressed SRF to drive arrector pili muscle maturation (Fig. 6J).

To investigate the similarity between maturing arrector pili muscle and fetal heart cells, we performed a cell-type signature enrichment across different developmental muscle cells, as well as the SkO and adult arrector pili muscle (Methods). Out of all cell types in the human body, this analysis showed enrichment for gene sets observed in prenatal heart, in prenatal skin, and SkO arrector pili muscle, suggesting that arrector pili muscle maturation shares components of the transcriptional program used during fetal cardiac development (Fig. S15e). The combination of associated transcription factors in the arrector pili muscle shows a similarity of arrector pili muscle to cardiac development (*MYOCD, KLF13, TCF21*) but also smooth muscle development (*SRF, MYOCD, MEF2A/C*). Thus, arrector pili muscle maturation appears to deploy a hybrid regulatory program that combines transcription factors characteristic of both cardiac and smooth muscle development.

### Sebaceous Gland Development is Characterized by Lipid Production and Cell Cycle Arrest

The sebaceous gland develops above the hair bulge and is responsible for sebum production through holocrine secretion (*136*, *137*) (Fig. 6A). In mice, the first glandular protrusions are visible around HF Stage 8, when the hair canal and IRS have formed (*39*). Whether the sebaceous gland is replenished by the HFSC populations in and around (*40*, *138*) the bulge or has a distinct stem cell population (*6*, *59*, *139*) has been debated. We found the earliest progenitors of the bulge (Stem-like PreHFSC) and sebaceous gland (PreSebocyte) to be located at the intersection of the ORS segments (Fig. 4G). Both progenitor populations are transcriptionally similar to the ORS segment they are found in (Fig. S16a). This lineage acquisition of PreHFSC and PreSebocytes before the mature structure has formed, together with the transcriptional similarity to the ORS segments, supports early lineage segregation within the developing ORS, consistent with the establishment of distinct stem cell populations.

To identify signaling factors that may drive the lineage segregation of PreSebocyte and Sebocyte, we applied MISTy human Stage 6 HFs (Methods). In the environment of PreSebocytes, the expression of *PTHLH* was the strongest predictor of PreSebocyte identity, followed by *ADAMTS1* and *LGR6* (Fig. S16b). Given this association, we asked whether altered *PTHLH* signaling prevents mature sebaceous gland formation in SkO HFs (Fig. 1D). *PTHLH* is initially expressed similarly to in vivo in the ORS infundibulum of Stage 6 SkO HFs (Fig. S13b), but is lost by day 130 (Fig. S16c), coinciding with the emergence of the aberrant KC: ORS Upper cell state. This KC: ORS Upper expressed stem cell markers such as *LHX2* and *LGR5* but no common ORS infundibulum markers (Fig. S16d). Investigating pathway activity, we identified enrichment of hypoxia genes in KC: ORS Upper (Fig. S16d,e), suggesting that hypoxia might be a driver of the aberrant ORS without a mature sebaceous gland in the SkO.

Investigating the drivers of maturation of human scalp PreSebocytes into mature Sebocytes, we identify the expression of many fat- and cholesterol-related metabolism genes, such as *SOAT1*, *GAL*, *PDZK1,* and *DHCR24* (Fig. 6B, C, S16f). We performed Hallmark (*140*) and CollecTRI (*141*) footprint-based TF enrichment analysis along the developmental process using a pseudotime with the infundibular keratinocytes as the starting point (Fig. S17a, Methods). While pathway enrichment showed a positive association of the developmental trajectory with *MTORC1* signaling, cholesterol, fatty acid, and bile acid metabolism, pathways that have been reported in adipocytes (*142*, *143*) (Fig. S17b), the regulon analysis demonstrated that *MTORC1* downstream genes Srebp-1 (*SREBF1*) and Srebp-2 (*SREBF2*), among Liver-X-receptors (*NR1H3*) and *PPARA/G*, are highly active in Sebocytes (Fig. 6D). While Srebp-1 is known to be a master regulator of lipogenesis in adult murine Sebocytes (*144*), Srebp-2 is known to regulate cholesterol production in the liver (*145–147*).

We compared our observed transcriptional program to the human adult sebaceous gland, where sebum is a minor product (*148*) (Methods) and found similar active TFs in the transition from Basal ORS to Basal Sebocytes and Sebocytes, with activation of *PPARA/G* and *SREBF1/2* (Fig. S17c). The prenatal scalp sebaceous gland had higher expression of *SREBF2* than adult ones (Fig. 5E).

To provide computational support for the role of prenatal sebaceous glands in cholesterol production, we predicted metabolite production in Sebocytes by metabolic enzyme enrichment analysis using a ULM with the producing and degrading enzymes of a metabolite from MetalinksDB (*149*) as weights (Methods). This analysis showed enrichment for Lathosterol-related genes in sebaceous glands (Fig. S17d), a cholesterol precursor reported as a reliable surrogate of cholesterol production in the body (*150*). These findings support the hypothesis that developing Sebocytes produce cholesterol. Increased cholesterol production may also be due to the demand for cell membrane generation during active prenatal development. Surprisingly, prenatal sebaceous gland cells have lower proliferation rates both in the infundibulum and in sebocytes (Fig. 6F) and have a smaller cell area compared to the adult sebaceous gland (Fig. 6G). This argues against cholesterol production to support membrane production. A potential use for the produced cholesterol could be the vernix caseosa, a white, high-cholesterol (*151*) lipid shell that surrounds fetuses *in utero,* preventing water loss and providing lubrication for the birth process (*152*) (Fig. S17e).

Similar to adipocytes (*153*, *154*), terminal sebocyte differentiation, including maximal lipid accumulation and holocrine secretion, is coupled to withdrawal from the cell cycle (*136*). Our data support this observation by showing an increase in G1-phase cells in Sebocytes compared with ORS Infundibulum and PreSebocytes (Fig. 6G, S17f). Sebocytes expressed high levels of CDKN1A (p21) while downregulating the G1 cyclins CCND1 and CCND2 (Fig. 6H, S17f), accompanied by increased chromatin accessibility at the CDKN1A promoter indicative of transcriptional activation (Fig. S17g). This suggests CDKN1A as a candidate regulator of sebocyte cell-cycle exit, similar to how it supports adipocyte hypertrophy in mice (*155*).

### Overriding of Prenatal Sebaceous Gland Gene Programmes in Sebaceous Tumors

The described spatial multiomic atlas provides a powerful lens to study skin disorders. Having characterized the transcriptional switch that halts the cell cycle and upregulates lipid production in developing and adult sebaceous glands, we investigated whether the transcriptional programme driving this switch is involved in sebaceous tumors, which can be benign (sebaceous adenoma and sebaceoma) or malignant (sebaceous carcinoma). We generated spatial transcriptomic data from these three tumor subtypes (n=3 per type, tissue collected in a previous study (*19*)) and processed them similarly to the scalp and SkO datasets (Methods). Tumour and non-tumoral Sebocytes clustered independently and aligned with clinical histopathology identification by H&E staining (Fig. 6I, S17h). We took the top 20 genes correlated and anticorrelated with sebaceous gland development, which contain mostly metabolism-related genes and cell cycle inhibitors, and defined them as the sebaceous gland developmental program. We then measured the enrichment of this gene program in the tumoral and non-tumoral Sebocytes using a univariate linear model (ULM) (Methods). We observed a lower enrichment in tumoral Sebocytes, which was significant in sebaceous carcinoma and sebaceoma (Fig. 6J; WLS regression, Benjamini–Hochberg FDR-adjusted *p* = 1.03 × 10^-4^ and 8.21 × 10^-4^, respectively). Lower enrichment of the Sebocyte developmental program was associated with a higher fraction of cycling cells (Fig. 6K). Consistently, sebaceous adenoma and sebaceoma retained high expression of lipid production genes such as PPARG (Fig. 6L) and Sebocyte-associated metabolic pathways (Fig. 6M). Sebaceous carcinoma notably showed diminished expression of p21, the driver of cell cycle arrest during prenatal Sebocyte development, as well as expression of cyclins and cyclin-dependent kinases (Fig. S17i) previously reported to be involved in sebaceous tumor development (*156*). p21 is also a downstream target of *NOTCH1*, which carries the most observed driver mutations of sebaceous tumors (*19*). Taken together, this suggests that sebaceous tumors override Sebocyte developmental programmes to become more proliferative, produce lower amounts of lipids, and the extent to which the developmental programme is lost tracks with malignancy (Fig. 6N). Although cancer evolution is generally associated with cellular dedifferentiation through the hijacking of developmental programmes, our findings suggest the opposite in the progression of sebaceous gland-associated tumours.

## Discussion

Here we apply a multimodal approach including spatial genomics at subcellular resolution to interrogate the full development of the human pilosebaceous unit, linking function to tissue geometry. We provide the most comprehensive classification of human prenatal scalp and iPSC-derived SkO hair follicle development, including the formation of the dermal sheath and arrector pili muscle for the first time. Furthermore, we uncover the stem cell/progenitor identity of the lower ORS cells, which are transient from the bulge HFSC compartment and replenish the proliferative matrix cells that produce the hair fiber. During the process, we find the lower dermal sheath and dermal papilla stalk act as a stem cell niche regulator, before WNT signals from the dermal papilla activate those cells to become proliferative. Our findings indicate that epithelial-mesenchymal interactions not only initiate the first stage of HF formation, but also orchestrate the entire HF formation and its subsequent maintenance. Finally, we describe how the sebaceous gland develops at the intersection of two spatial segments and how this physiological developmental process is overridden in sebaceous tumors.

The main advantage of our study is the use of high-quality, high-plex spatial transcriptomics coupled with H&E staining of the same tissue section and multiome data from the same tissue blocks. The integration of spatially resolved molecular profiles at single-cell resolution with known HF tissue morphology allowed us to map small differences in the transcriptome to the pilosebaceous functional units to derive fine-grained annotations. An example of this is the detection of the EDC and dermal sheath. This previously unknown EDC population is condensed around placodes, transcriptionally distinct and SOX2-negative, in contrast to the murine pre-DC (*63*, *157*). Recent studies on murine placode development have found a ring of fibroblasts that support placode formation by contracting (*58*). This contraction is exerted by myosin 2 light and heavy chains, both of which function in conjunction with MYLK, which was also demonstrated to be a driver of contraction in the adult dermal sheath (*71*). Given that *MYLK* is a key marker of the EDC we observed, it is intriguing to speculate if this early population has a contractile phenotype that helps the downgrowth of the follicle.

In addition to its location around the early placodes, we could show that the EDC gives rise to the proliferative EDS through being exposed by the tissue geometry to signaling gradients of the downgrowing hair peg. While being identified as an *SLC26A7*+ “fibroblast hair” population in a recent study (*70*), it was often not included in many previous studies either because it was not captured (SOX2-)(*63*, *157*) or labelled as dermal condensate (*47*, *157*). In agreement with our study, murine Dkk1 is expressed in the EDS, but not the dermal condensate, and loss of Dkk1 arrests hair follicle formation (*30*). The hypothesis of dermal sheath descendancy from dermal condensate progenitors rather than dermal fibroblasts is supported by lineage tracing of Tbx18+ (*72*) and Foxd1+ (*158*) dermal condensate cell descendants in mice. The identification of the molecular profile of the EDS population might allow their isolation or directed differentiation in vivo in the future, which holds therapeutic value, as the dermal sheath contains stem cell properties and is involved in wound healing.

Another strength of the spatial data is to detect molecular patterns that are not visible in histology, an aspect recently discussed in the context of pathological immune memory niches around adult sebaceous glands (*52*). An example is the two transcriptionally distinct but histomorphologically identical segments of the ORS that determine the positioning of the HFSC and sebaceous gland. These segments are also observed in middle-stage hair in SkOs, but are not maintained throughout the HF developmental stages in vitro.

The link between form and function also manifests in the final form of the pilosebaceous unit. This principle is demonstrated by the geometry of the bulge and bulb that together support hair shaft production and regeneration in an opposing “escalator-like” movement between the ORS cells downwards and the hair shaft upwards. Through identifying the stem-cell identity of the lower ORS, we reveal a link between the dormant HFSC reservoir in the bulge that moves downwards to replenish the proliferative KC: Matrix in the bulb and gives rise to the hair shaft and the IRS, which move superficially upward. While the downward movement of the ORS has been reported (*94*, *95*), we show their stem cell/progenitor molecular profile for the first time. We further show global coordination between the lower ORS and the adjacent mesenchymal dermal sheath that supports the HFSC of the lower ORS (*105*) staying dormant, extending the known role of dermal sheath beyond biomechanical functions in the hair cycle (*71*). This finding emphasizes how the hair form optimizes the interface for epithelial-mesenchymal interactions during hair development and maintenance.

Another example of global coordination is the development of the arrector pili muscle. Given that piloerection is unlikely to be required *in utero*, we expect the arrector pili muscle to be functional only after birth. Accordingly, we find that mature lanugo hair is associated with an immature arrector pili muscle, suggesting that maturation follows functional requirements rather than anatomical completion. As the arrector pili muscle represents a highly specialized smooth muscle subtype, its maturation provides a model for understanding how smooth muscle diversity is established during development. We show that muscle maturation is characterized by MYOCD and SRF activity, both known to be regulators of several muscle types, supporting the idea that combinations of shared transcriptional regulators contribute to muscle subtype specification.

In addition to these biological findings, we use the decision-tree-based method XGBoost under the MISTy framework to study tissue microenvironmental drivers of differentiation processes. Using the spatial information of the cells to construct a temporal ordering, and leveraging the interpretability of the XGBoost model, we can predict tissue microenvironmental features that lead to bifurcation points in spatiotemporal trajectories, such as the differentiation of the dermal sheath and condensate. This use of spatial information for ground truth trajectories was recently demonstrated (*74*) and will allow many more approaches to go from steady-state inferences to dynamic analysis of system behavior. Furthermore, this manual temporal ordering will have resource value as a validation dataset for perturbation prediction algorithms.

The data presented in this study are mostly observational and thus come with technical limitations. Our analysis highlights many predicted progeny relationships, such as the first progenitors of dermal sheath, CL, or IRS, which are based on transcriptional and epigenetic similarity as well as spatial observations. While equivalent experiments have often been done in mice (*57*, *72*, *158*), suitable follow-ups would include lineage tracing experiments, for example, barcode-based in SkOs or via tracing of methylation and somatic mutation patterns *in vivo*. Another limitation originates from the segmentation before cell calling, which can be challenging in densely populated areas such as keratinocyte layers or cells with elongated shapes such as neurons. Common segmentation errors can lead to transcriptomes acquiring characteristics of the surrounding cells, which was mitigated by giving lower weight to analysis results including cell types with high expected non-self contribution in their transcriptome, such as melanocytes.

In conclusion, this work is the first cell-type-resolution spatial map of a full mini-organ development that shows how, in a conserved structure such as the pilosebaceous unit, constituent components interact with each other to guide morphogenesis and shape a geometry that allows multiple overlapping functions to be executed at the same time. Additionally, this study shows how different experimental and computational approaches can be combined to interrogate these spatiotemporal signaling patterns. These insights will feed into studies inducing hair *de novo*, and hold potential to improve pre-clinical models, skin engineering and hair renewal.

## Acknowledgements

We thank Aidan Maartens, Fränze Progratzky, Jovan Tanevski, Pierre Murat, Vicky Moignard, Pasha Mazin, John Lee, Ciro Ramírez-Suástegui, Koen Rademaker, Ricardo Ramirez Flores, Arran Constantine, Marc-Antoine Jacques, Francesco Carli, Adam Boxall, Masami Ando Kuri, Anna Pournara, Kwasi Kwakwa, Francesca Drummer, Denes Turei, Phillip Sven Lars Schäfer, Charlotte Boys, and Benjamin Rumney for helpful discussions. Biorender was used for select figure design. Generative AI was used to improve code and text clarity.

## Author contributions

Conceptualization: AF, MH, EF

Data curation: AF

Data generation: AF, EK, MC, CA, ES, KE, MP, FT, HMC, CT

Formal analysis: EF

Funding acquisition: MH, DJA, JSR

Investigation: EF, MH, AF

Methodology: EF, AF, AD, MH

Project administration: AF, EF, MH

Resources: AF, DH, MPr, LG, KS, DA

Software: EF, VB, DH, JB, AP, AB, TL, DBL, KP

Supervision: AF, AD, MH, VB, JSR, LP, DJA, ES

Validation: AF, EG, EF, JM, VB

Visualization: AF, EF, DBL

Writing – original draft: EF, MH

Writing – review & editing: MH, AF, HG, LS, EW, MK, CS, JM, LP, BH, DJA, FT, CH

## Supplementary information

Material and Methods

Figs. S1 to S17

Tables S1 to S8

References 159-175

