## Supplementary text and figures for "Function-driven geometry directs human pilosebaceous unit development"

### Material and Methods

#### Ethics for tissue collection

All prenatal samples were collected under the MRC–Wellcome Trust-funded Human Developmental Biology Resource (http://www.hdbr.org) with approval from the Newcastle and North Tyneside NHS Health Authority Joint Ethics Committee (08/H0906/21+5) and East of England–Cambridge Central Research Ethics Committee (NHS REC 96/085). Healthy adult trunk skin samples were collected with full approval from the Newcastle and North Tyneside NHS Health Authority Joint Ethics Committee (NHS REC 24/NE/0014). Sebaceous tumor samples were collected as part of a larger study that has been approved by the National Health Service Health Research Authority (Research Ethics Committee reference 21/PR/1024, IRAS project ID 304621).

#### Tissue collection and processing

For prenatal scalp skin samples, tissue was expertly dissected and embedded in optimal cutting temperature compound (OCT). Samples were sectioned at 10 um thickness on a cryostat with an effort to align the block with the growth angle of the HF for profiling via spatial transcriptomics. To process skin samples for Multiome, eight 25 μm sections were collected into chilled Eppendorf tubes and then taken through the protocol for 10X Multiome data generation following the manufacturers' protocols. For adult skin samples, skin biopsies were collected and formalin-fixed, then paraffin-embedded. Samples were sectioned at 5 μm thickness on a microtome for performing spatial transcriptomics. The sebaceous tumor data were obtained from samples acquired in a previous study (*19*).

#### Processing of skin organoids

SkO culturing was performed as described in this study (*46*). Macrophages and myeloid progenitors were added at day 12 of SkO as described in our previous study (*47*). Whole-mount organoids were cultured from day 63 in air-liquid interface culture using a transwell system (Corning). For imaging spatial transcriptomics, SkOs were embedded in OCT at day 34, 54, 76, 95 and 130, and sectioned at 10 um thickness on a cryostat.

#### Xenium Panel design

We designed a custom 100-gene panel (Table S8) using the Xenium panel builder. Genes were selected from the literature and in-house datasets. Additionally, we added genes that were known to be either cell type markers or genes with a high spatial autocorrelation (Moran's I) or HOX genes in various in-house datasets.

#### Preprocessing of multiome and spatial data

Cells were called using the Xenium standard segmentation. For the developmental dataset, we excluded cells with fewer than 100 unique genes detected. One section (Posterior, PCW19) was excluded due to tissue detachment. To exclude folded or overlapping regions, we drew polygons in Xenium Explorer v4.0.0 and excluded all cells outside those polygons (Fig. S1). Subsequently, to allow visual inspection of all sections at once, their coordinate systems were rotated to have the epidermis up and spread in a way that the x-axis described the time in post-conception weeks and the y-axis described from which scalp region the samples were taken (Fig. S1).

For the SkO Xenium data, no cells were excluded for a low number of genes detected, due to the suprabasal keratinocytes having a low number of total counts and being challenging to separate from low-quality cells when utilising quality control metrics alone. Cells lying outside the tissue sections were excluded using manually set xy thresholds. Afterwards, the coordinate systems of the sections were rotated, flipped, and aligned to allow interrogation of the different sections.

For the multiome, preprocessing was performed using cellranger-arc v2. The RNA modality was then filtered for high-quality cells with less than 60 % of counts in the top 100 genes, 15 percent ribosomal counts, 10 % hemoglobin, 2 % mitochondrial counts, and a Scrublet(*159*) doublet score lower than 0.1.

The ATAC data was processed with a pycistopic (*160*)-based Nextflow pipeline (https://github.com/cellgeni/nf-atac, release 26-079) that takes the ATAC fragments and cell type labels as input. It outputs bigwig files for visualization, pycistopic, and h5ad objects that are used for further downstream analysis.

#### Integration of datasets

Xenium and multiome datasets were integrated using a custom pipeline that compares several integration and feature selection methods as well as different numbers of highly variable genes using scIB (*161*). We decided on 4500 highly variable genes called by the “cellranger” option in scanpy v1.10.0 (*162*) using the sample as batch key and scVI v0.6.8 (*49*) with 2 layers, based on in-house benchmarking of several combinations.

#### Annotation

Before the annotation, a hierarchical classification of cell types at several resolutions was constructed. Following that, cell types were classified top-down, starting with the biggest groups (epithelial, dermal stroma, musculoskeletal, etc) to the smallest ones (hair shaft Inner, melanocyte HF, etc.). Classification usually followed the scheme of clustering parts of the dataset and then looking at the differentially expressed genes (see section differential abundance testing) and the spatial location of the clusters. The full preprocessing and annotation notebooks will be available on github. A table with markers per cell type can be found in Table S1.

To annotate the RNA of the multiome data, the multiome and Xenium objects were concatenated on their shared genes and integrated using the above-mentioned pipeline. The labels of the Xenium were then transferred to the multiome using a custom KNN classifier on a KNN graph constructed on the integrated scVI (*49*) latent space (github). In brief, cells were assigned their neighbors' labels if there was a clear majority of labels. In the first round, all cells were assigned if more than 80 percent of the nearest neighbors had the same label; in the next round, 70 percent, and so on, going down with the percentage criterion until all cells were labelled. This process allows regions with few labelled nearest neighbors to first get a few good confidence labels, before they are assigned to a cell type based on very few cells. The validity of the label transfer was determined by measuring per-gene (Fig. S1e) and per-cell-type (Fig. S1f) correlation between the corresponding labels of the two data types.

#### Inference of transcription factor, ligand, pathway, and cell cycle activities

TF activity analysis was performed using a ULM available in decoupler (*163*) on literature-curated downstream genes of TFs (*141*), where genes that are known to be activated by a certain TF get a value of 1, and genes that are known to be downregulated get a value of -1. NicheNet (*164*)-like ligand activity analysis was performed, similar to the TF activities, using the ligand target matrix from NicheNet v2 (*165*) (ligand_target_matrix_nsga2r_final.rds, https://zenodo.org/records/7074291). Reactome (*166*), cell type signature (MSigDB (*167*) C8), and Hallmark (*140*) analysis of the respective gene sets available in Omnipath (*168*) using a ULM in decoupler (*163*) as above. The cell cycle stage was inferred using the Scanpy "score_genes_cell_cycle" function with genes found on GitHub (https://github.com/scverse/scanpy_usage/blob/master/180209_cell_cycle/data/regev_lab_cell_cycle_genes.txt).

##### Inference of cell-cell communication

Dissolved cell-cell communication analysis was performed using the 'rank_aggregate' function in LIANA+ (*73*) that runs a total of 5 methods and aggregates the ranking of a co-expression LR pair across methods in a magnitude (analogous to, e.g., a fold-change value) and specificity (analogous to a p-value) rank score.

##### Differential abundance and expression testing

Differential abundance was tested using milopy (*61*) with default parameters.

Differential gene expression analysis was done using a Wilcoxon signed rank test in the scanpy "rank_genes_genes_groups" function unless specified otherwise.

##### Inference of expression trajectories

Expression trajectories of selected cell types were inferred by dpt_pseudotime in scanpy, usually using anatomical locations (e.g., placodes) as starting points. As the next step, the pseudotime was checked for correlations with the cell cycle and absolute time to confirm validity or lower the likelihood of false interpretation. Relevant gene expression or transcription factor activity patterns along pseudotime were then determined using the "rank_order" function in decoupler.

##### Inference of metabolite abundance

Metabolic enrichment was performed using the module in LIANA+ (*73*), which runs a ULM using decoupler-py on a database of metabolic reactions called MetalinksDB (*149*).

#### Assembly of HF stages

For the assembly of HF stages, a total of 137 HFs from all stages were cropped and rotated to align along the y-axis. A table with coordinates and rotation angles can be found in Table S2. The cropped follicles were then staged and put in order. For Fig. 1D, representative samples were chosen to reflect the whole span.

#### Determination of gradients along tissue depth

To determine which genes were expressed with gradients concerning the distance to the epidermis, cropped, rotated HFs (see section assembly of HF stages) from the relevant stages were selected. From these follicles, the gene expression of the outer and IRS cells was correlated with the y-position of the selected cells using the Spearman correlation of the decoupler correlate_order function. Similarly, the transcription factor and ligand activity gradients were determined. For the tissue gradients in Fig. S8c, all relevant cells were correlated with the y-axis.

#### ATAC analysis

Consensus peaks were filtered for high confidence by excluding peaks with a normalized accessibility of 2. Afterwards, co-accessible peaks were determined using circe (*169*), a fast Python-based analogon to cicero (*170*). Peaks were mapped to transcription start sites, and TF motif binding was performed using CellOracle (*133*).

#### Co-occurrence analysis

Co-occurrence of cell types was determined using the “compute_co_occurrence” function in squidpy(*171*) v1.6.2, with a tiling allowing bins of approximately 10 um spanning a distance from 0 to 100 um with 10 um steps in between.

#### MISTy analysis of temporally ordered HF

To obtain a temporal ordering, the coordinate systems of cropped HF (see above) were aligned with the y-axis. We then measured the distance from the epidermis to the distal tip of the hair follicle and ordered them after increasing distance. XGBoost runs on two classes, which were assigned based on I) gene expression for EDC and IRS predictions and II) spatial location for EDS/DC, PreHFSC, and PreSebocyte prediction. The two classes are then one-hot encoded as the intraview and the expression of all genes in the surrounding cellular environment as the paraview using a Gaussian kernel with a bandwidth of 200 um and a gamma of 30 um, roughly matching the diffusion radius of cytokines (*172*). We reduced the features using a threshold based on the product of spatial autocorrelation (Moran's I) and conditional gene expression variance. For the prediction, we used XGBoost (*173*) implemented in Scikit-learn (*174*) using the following parameters: a maximum tree depth of 3, 100 boosting iterations, a learning rate of 0.1, a subsampling ratio of 0.8, a column subsampling ratio of 0.8 per tree, a minimum child weight of 1, the binary logistic objective function, and log loss as the evaluation metric parameters. Cells were grouped based on full structures, e.g., placodes or hair shafts that had balanced class distribution, and split 80:20 into train and test sets. Predictions were repeated 3 times with different 80:20 splits and averaged afterwards. Feature importance was extracted using Shapley values (*175*) implemented in the SHAP Python package.

#### Immunohistochemistry

Immunohistochemical staining was performed as reported previously (*47*). In brief, slides stored at −80°C are thawed and allowed to dry at room temperature (RT). Dried slides are fixed in 4% paraformaldehyde (PFA; Thermo Fisher Scientific) for 10 min at RT in a chemical fume hood, then rinsed in 1× PBS (Thermo Fisher Scientific) for several minutes. The staining chamber is hydrated by adding water to the bottom of the reservoir, and slides are mounted in the device.

Each slide is incubated with 120 µl of blocking solution (3% goat serum (Abcam) in PBS and 1x Triton (PerkinElmer)) for 1 h at RT. Without rinsing, 120 µl of primary antibody mix in blocking solution is added to each slide, except for negative control slides. The chamber is covered with aluminum foil to protect from light and incubated overnight at 4°C.

The following day, slides are washed three times with 1 ml of 1× PBS per slide. Slides are then incubated with 120 µl of secondary antibody mix in blocking solution for 1 h at RT, followed by three additional washes with 1 ml of 1× PBS per slide.

When the mounting medium does not contain DAPI, slides are incubated with DAPI (1 µg/ml; Thermo Fisher Scientific) for nuclear counterstaining, followed by two washes in 1× PBS. Slides are air-dried, mounted with Vectashield antifade mounting medium (Vector Laboratories) or equivalent, and allowed to dry overnight at RT before imaging.

#### Wholemount staining

Organoids cultured on transwell plates were fixed in 4% PFA overnight at 4 $^{O}$C, washed in PBS, and kept at 4 $^{O}$C until staining. To begin the whole-mount staining process, organoids were transferred into an 8-well IBIDI plate and incubated in 50% CUBIC-L (TCI chemicals, T3740, diluted in ddH$_2$0) for 6 hours at 37 $^{O}$C, prior to approximately 18 hours of incubation in 100% CUBIC-L at 37 $^{O}$C.

Following this, each well was washed while being gently agitated with PBS for two 15-minute washes, and one 30-minute wash, before incubating overnight at 37 $^{O}$C in permeabilisation buffer (PBS containing 0.2 % Triton-X, 306 mM glycine, 10 % DMSO).

On day 3, the permeabilisation buffer was replaced with blocking buffer (PBS containing 0.2 % Triton-X, 6 % goat serum, 10% DMSO) and incubated at 37 $^{O}$C for 7 hours. Primary antibodies were diluted to desired concentrations (ACTA2 *ab124964* 1:100, Myocardin *MAB4028* 1:100, KRT15 *sc-47697* 1:100) in primary antibody buffer (PBS containing 0.2% Tween-20, 10 µg/ml Heparin (STEMCELL 07980), 5 % DMSO, 3 % goat serum). These were incubated for approximately 64 hours at 4 $^{O}$C with gentle agitation.

The primary antibody mix was removed and washed with PTwH (PBS containing 0.2% Tween-20, 10 ¨g/ml Heparin (STEMCELL 07980)) for 6 hours, refreshing at least three times. Secondary antibodies were prepared in secondary antibody buffer (PBS containing 0.2% Tween-20, 10 µg/ml Heparin (STEMCELL 07980), 3 % goat serum) and filtered with a 0.2 µm filter unit before use. Samples were incubated overnight at 4 $^{O}$C with gentle agitation.

The following morning, samples were washed with PTwH (PBS containing 0.2% Tween-20, 10 ug/ml Heparin (STEMCELL 07980)) for 6 hours, refreshing at least three times. After washing, samples were incubated in 50% CUBIC-R (TCI chemicals, T3983, diluted in ddH$_2$0) for 2 hours at room temperature, before moving to 100% CUBIC-R+ for at least 24 hours at room temperature. Samples were then imaged using a Nikon Ti2 confocal in mounting solution (RI 1.520) (TCI chemicals, M3294).

#### Fig S1: Preprocessing and integration allow consistent annotation across datasets

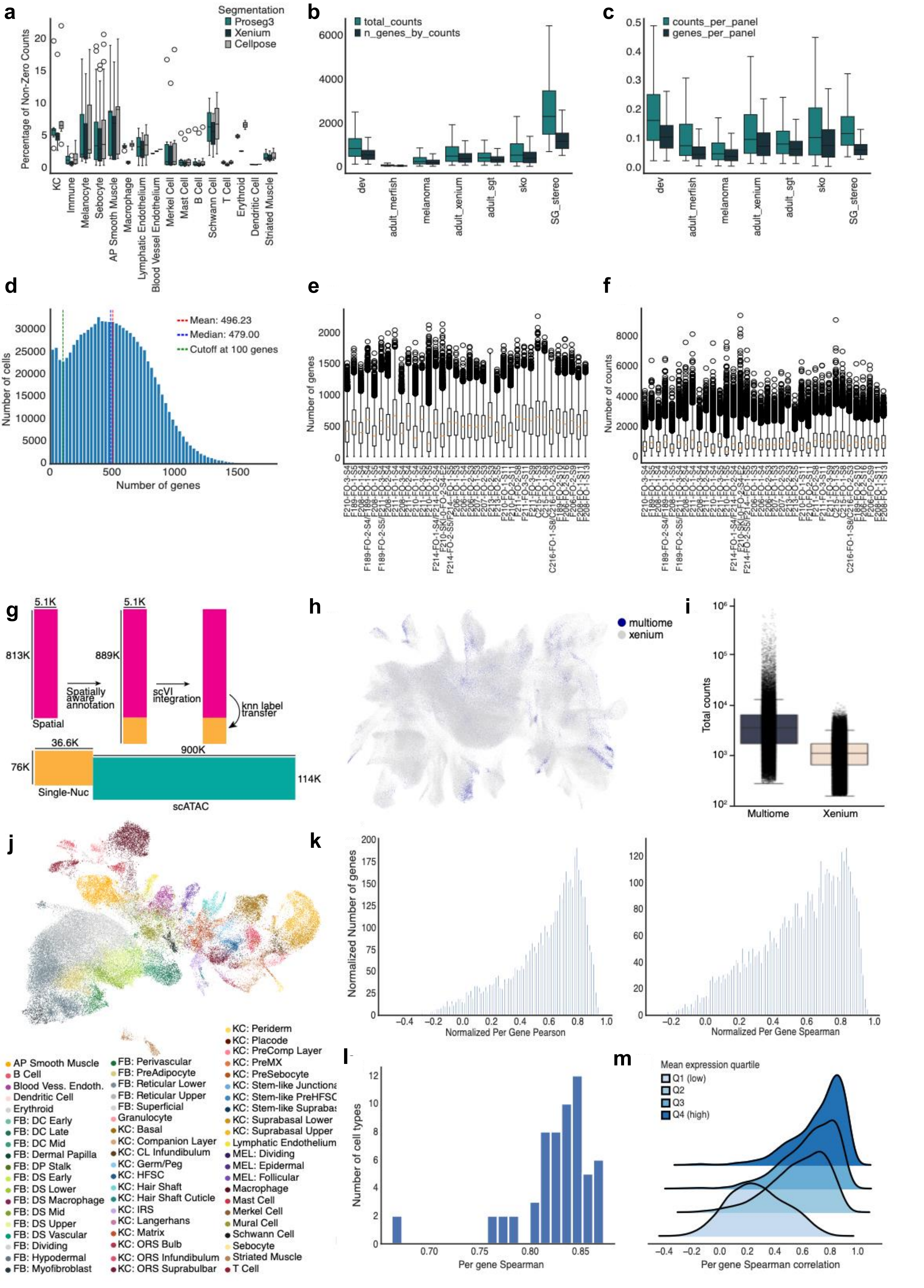

**Fig S1: Preprocessing and integration allow consistent annotation across datasets**

a Comparison of different segmentation methods by non-zero counts of cell type marker genes outside cell type. ProSeg3 and CellPose have consistently more non-zero counts indicating worse segmentation performance.

b Boxplot showing median genes and counts per cells of the fetal scalp as well as other published datasets. The fetal dataset has substantially more genes and counts then the published datasets.

c Boxplot showing median genes and counts per cells of the fetal scalp as well as other published datasets normalized by panel size. The fetal dataset has substantially more genes and counts then the published datasets.

d Distribution of numbers of genes of fetal scalp spatial data and cutoff line at 100.

e Number of genes per samples, shows stable medians.

f Number of counts per sample shows medians between 500 and 1500.

g Approach to map spatial transcriptomic to Multiome data. First the spatial data gets annotated and integrated on the same features then the Multiome RNA. In the shared latent space, the labels can be transferred via a KNN-classifier

h UMAP of shared latent space between spatial and Multiome RNA shows good mixing between the two datatypes

i Distribution of total counts between Multiome RNA and spatial show generally higher sensitivity of Multiome RNA than spatial xenium

j UMAP showing the mapped back labels from the label transfer. they show clear separation in the UMAP indication them explaining the Multiome RNA well

k Distribution of per gene Pearson and Spearman correlation between cell type level pseudo bulks from both datasets.

l Distribution of per cell type Pearson correlation between the cell type level pseudo bulks from both datasets

m Distribution of per gene correlation sorted after mean expression per gene, indication lower correlation of lowly expressed genes.

Fig S2: Annotation is driven by spatial annotation

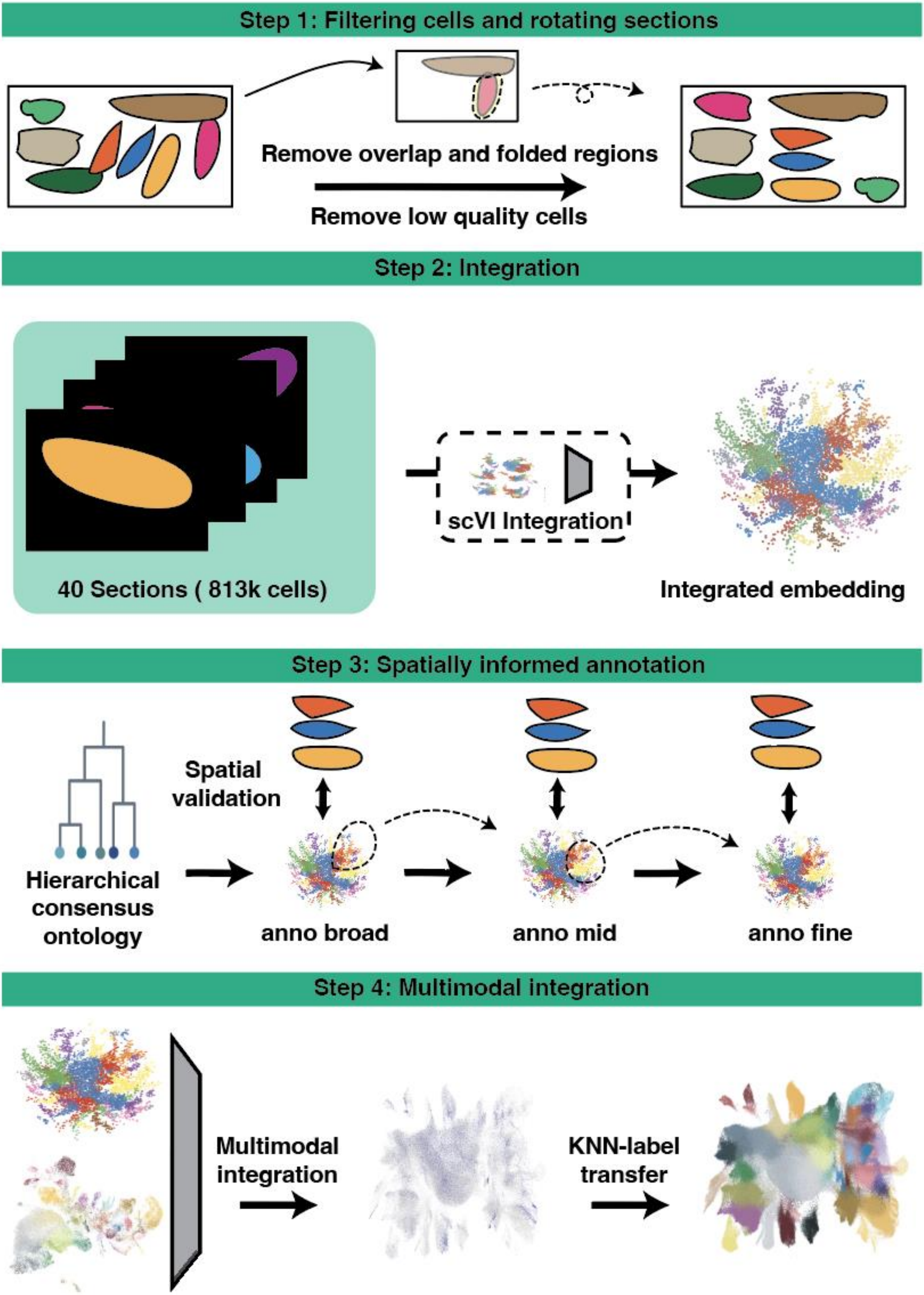

Fig S3: Spatial annotation reveals heterogeneity of spatial hair follicle compartments

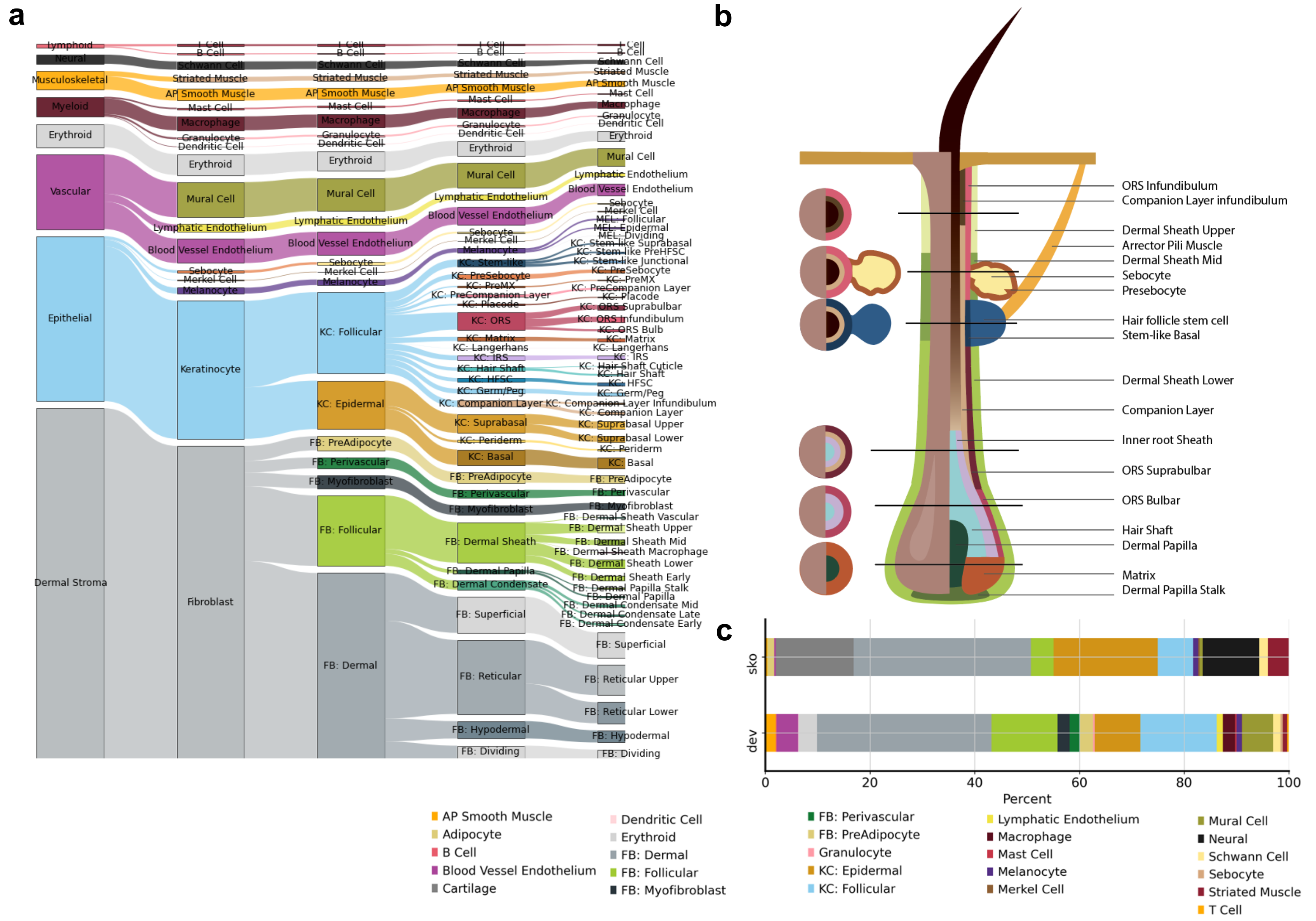


**Fig S3: Spatial annotation reveals heterogeneity of spatial hair follicle compartments**

a  Structure of annotation on 5 levels, with the level 5 having the highest granularity of cells with substantial annotations in the hair types.

b  Depiction of anatomical descriptions of developmental scalp hair.

c  Stacked bar plot showing granular cell type fractions between skin organoid and developmental skin.

### Fig S4: *In vivo* atlas shows location of immune cells

a

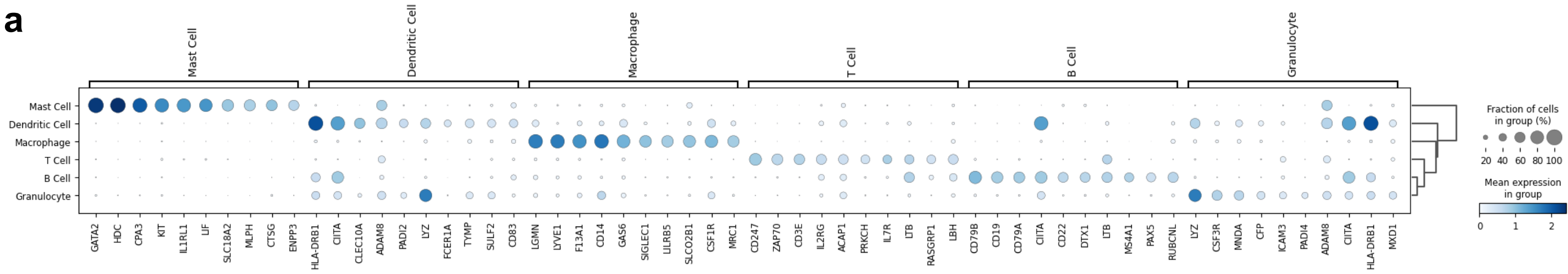


b

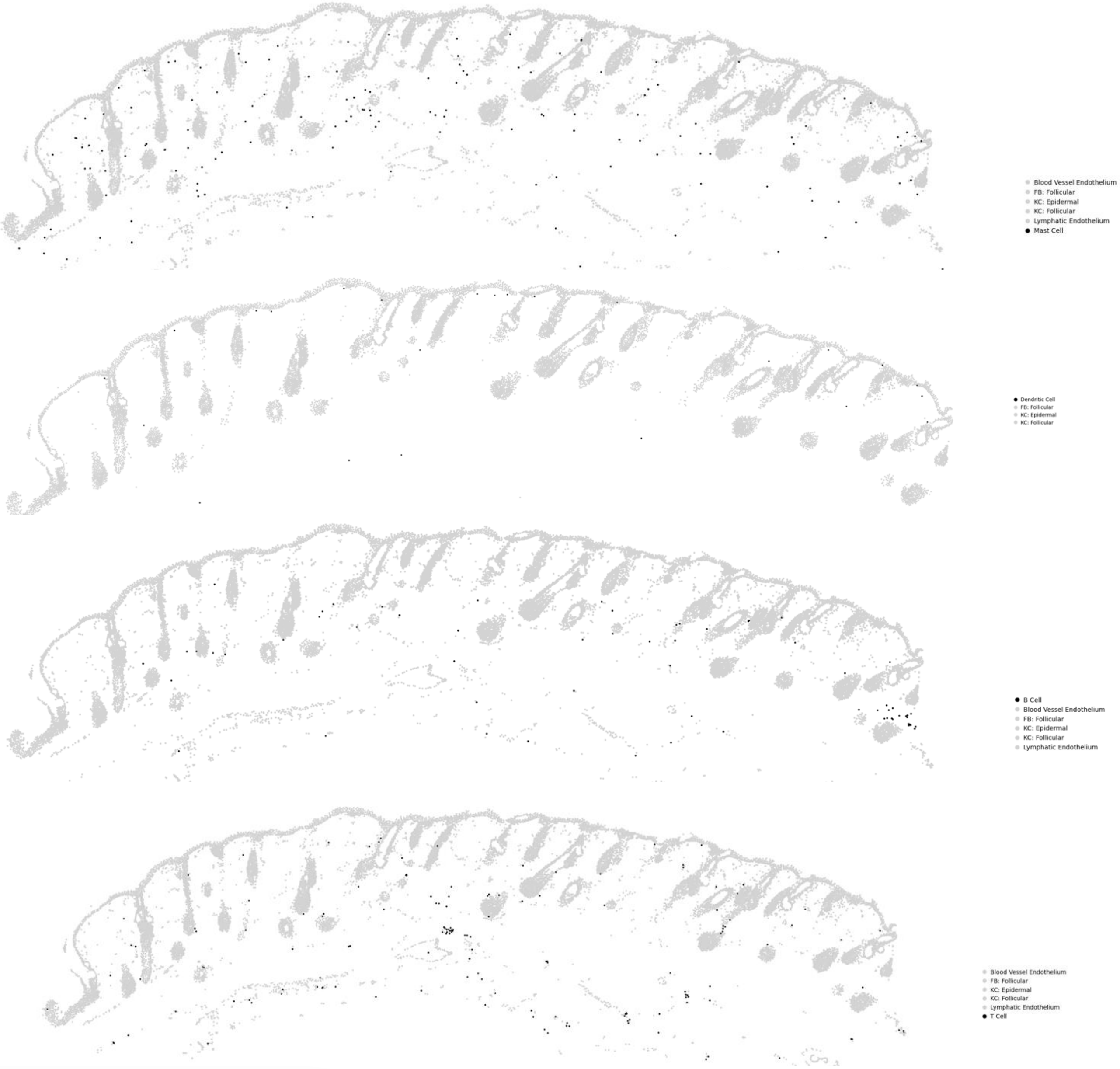


c

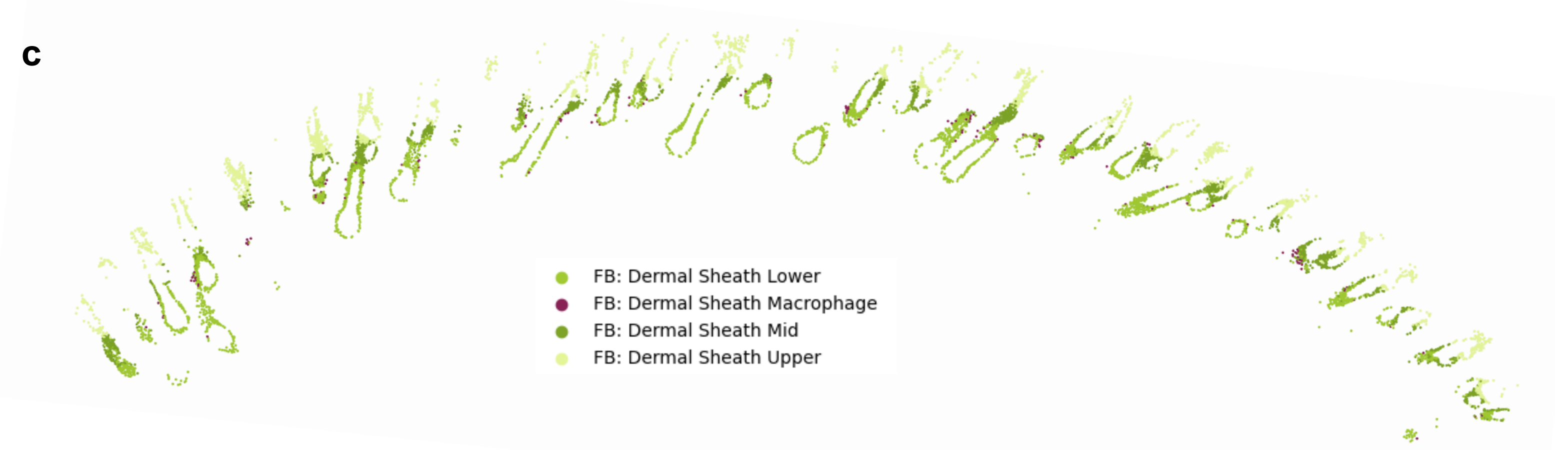


**Fig S4: *In vivo* atlas shows location of immune cells**

a Cell type markers of immune cells in developing scalp skin.

b Location of immune cells in the tissue and around vessels.

c Macrophage locate in the lower dermal sheath.

#### Fig S5: Placode formation can be observed across regions and gestational time

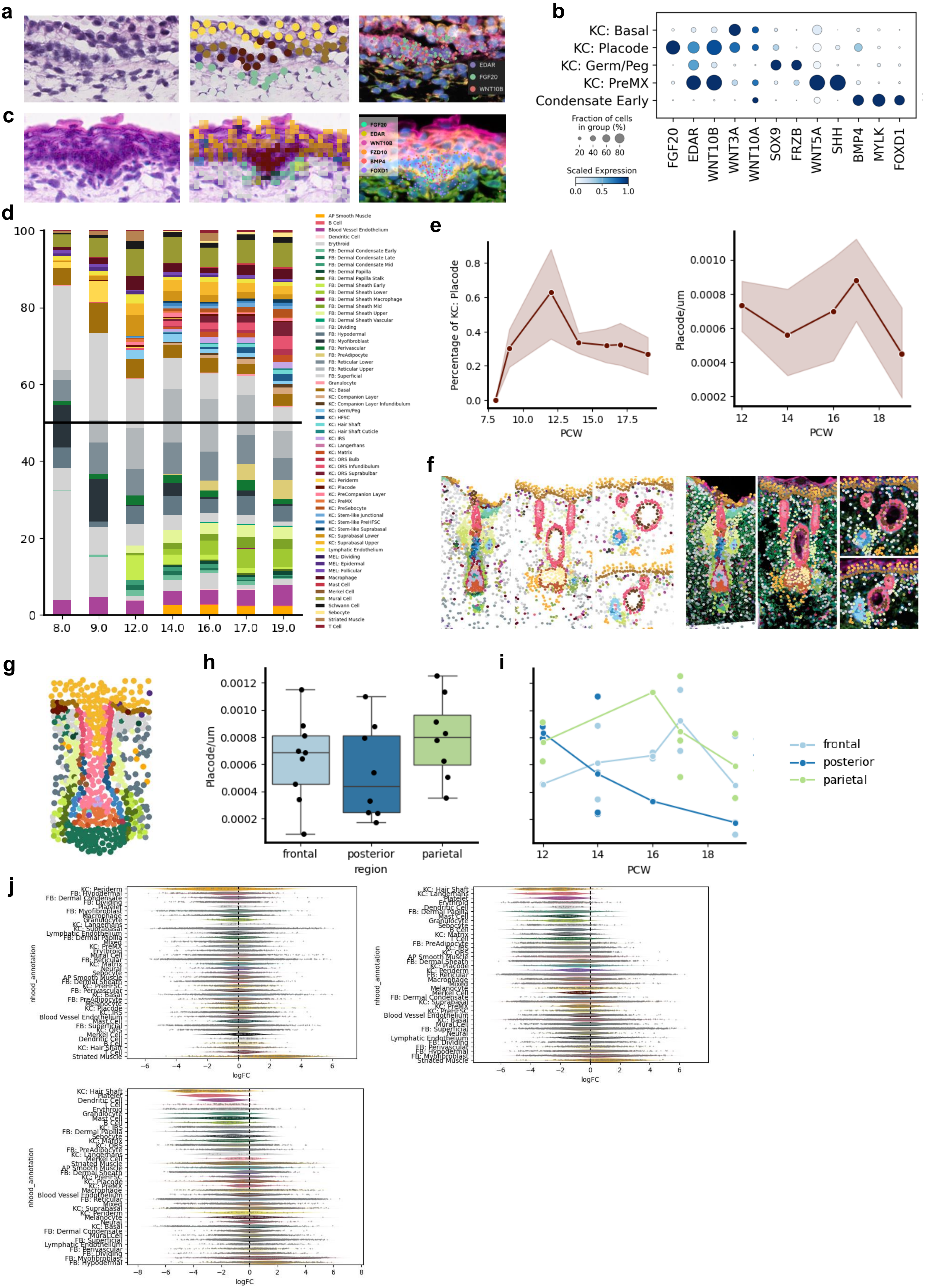

**Fig S5: Placode formation can be observed across regions and gestational time**

a Placode from PCW 9 without apical basal elongation of keratinocytes.

b Marker genes of early follicular keratinocytes.

c Placode from PCW 12 with apical basal elongation of keratinocytes.

d Stacked bar plot showing the cell type contribution over various post conception weeks.

e Left: Percentage of Placode cells per PCW, Right; Placodes per um of PCW 12 - 19, length of surface was measured by fitting lines to the basal keratinocyte (methods).

f Examples of placode first forming next to existing follicles (left) and triplets of hair with two young and an old hair with longitudinal (middle) and cross (right) section.

g Stage 6 Hair follicle from skin organoid.

h Placodes per um of PCW 12 - 19 between different scalp regions.

i Development of placodes per um over time show decrease of placodes per μm especially in the posterior region.

j Milo enrichment shows no substantial differences between regions.

### Fig S6: Early human hair mesenchyme has specific markers

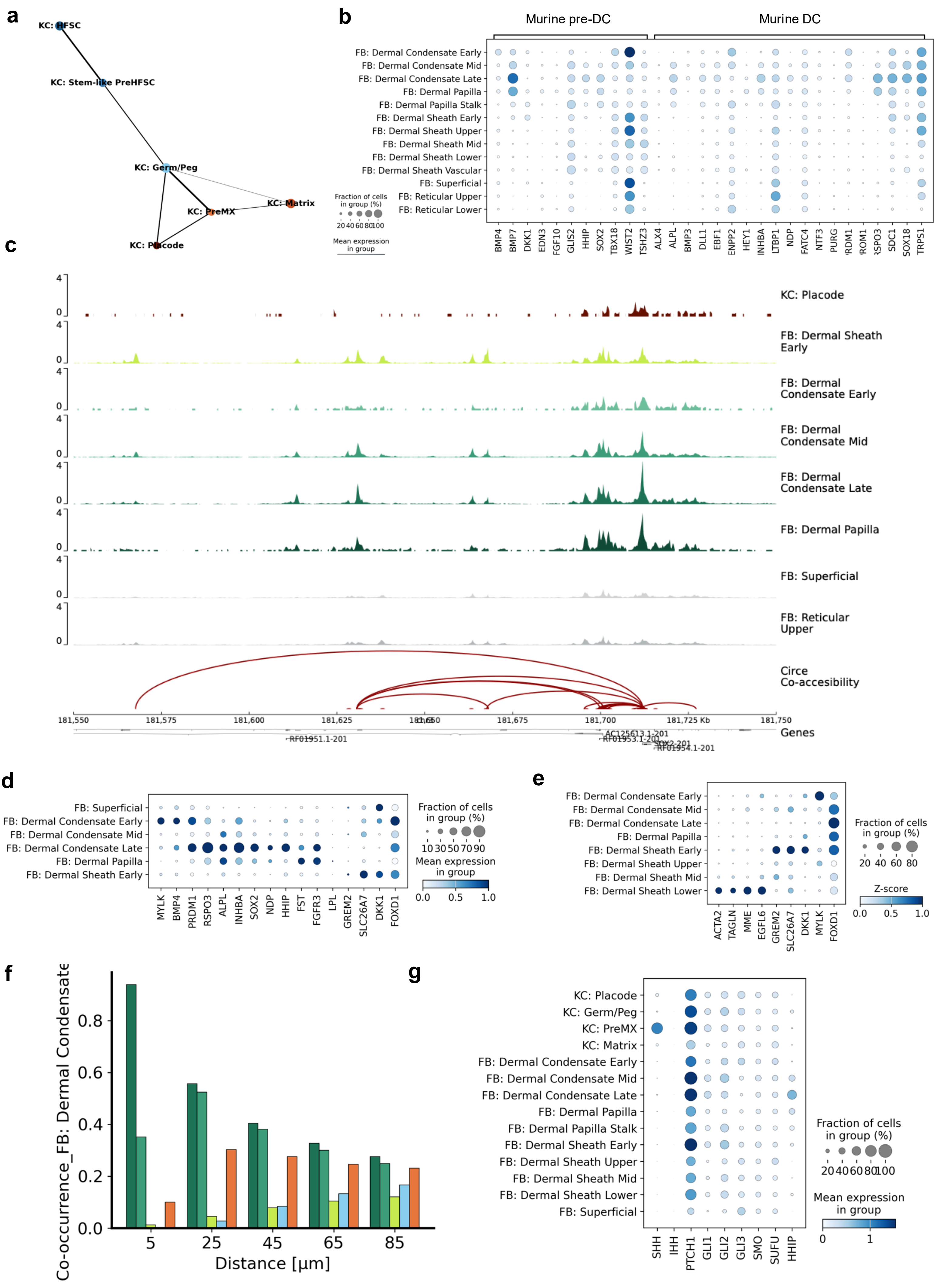

**Fig S6: Early human hair mesenchyme has specific markers**

a PAGA plot showing the similarity of cell types. The placode in the bottom develops into the PreMX and KC: Germ/Peg that themselves develop into HFSC and Matrix.

b Dot plot showing expression of murine pre-DC and DC marker genes.

c Chromatin accessibility of *SOX2* locus.

d Skin organoid markers or early and late dermal condensate.

e Markers of different dermal condensate and dermal sheath populations, showing that the smooth muscle phenotype is only acquired in the lower dermal sheath.

f Co-occurrence of early HF populations with EDS (FB: Dermal Sheath Early) 5-85 um

g Dot plot showing the gene expression of hedgehog pathway genes across all early hair follicle populations. SHH is only expressed in PreMX cells, while HHIP a negative feedback regulator of SHH is strongest expressed in dermal condensate late.

**Fig S7: Spatially informed temporal ordering allows identification of bifurcation point and its environmental drivers**

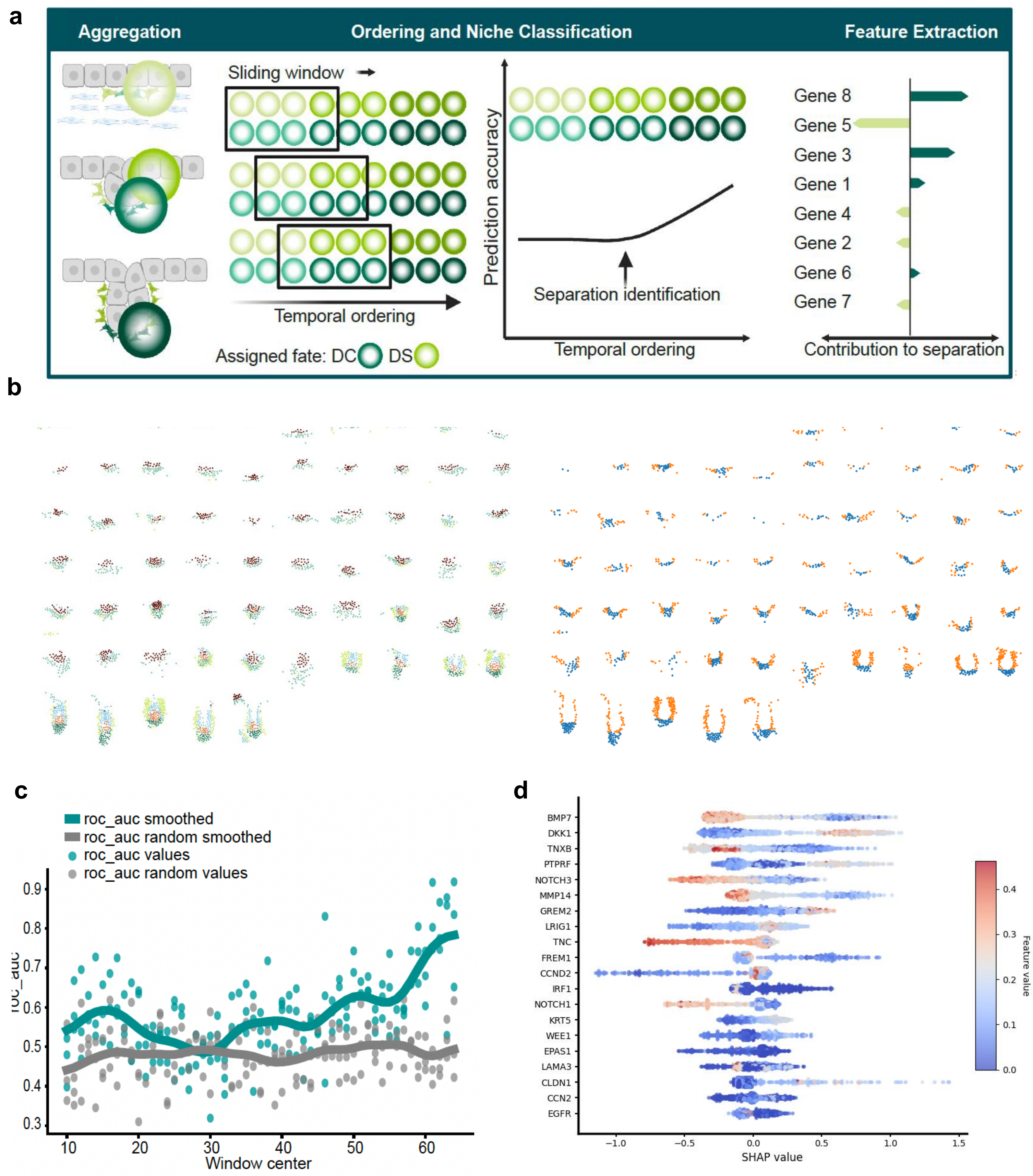


**Fig S7: Spatially informed temporal ordering allows identification of bifurcation point and its environmental drivers**

a Cartoon showing the principle of ignite in 4 steps: classification, aggregation, prediction and explanation.

b Placode ordered after age and coloured by gene annotation and class assignment.

c Result of sliding window approach predicting environmental factors with increasing prediction power.

d Features with the top Shapley values of the predictive model.

### Fig S8: Adipocytes form around hair bulb in lower reticular dermis

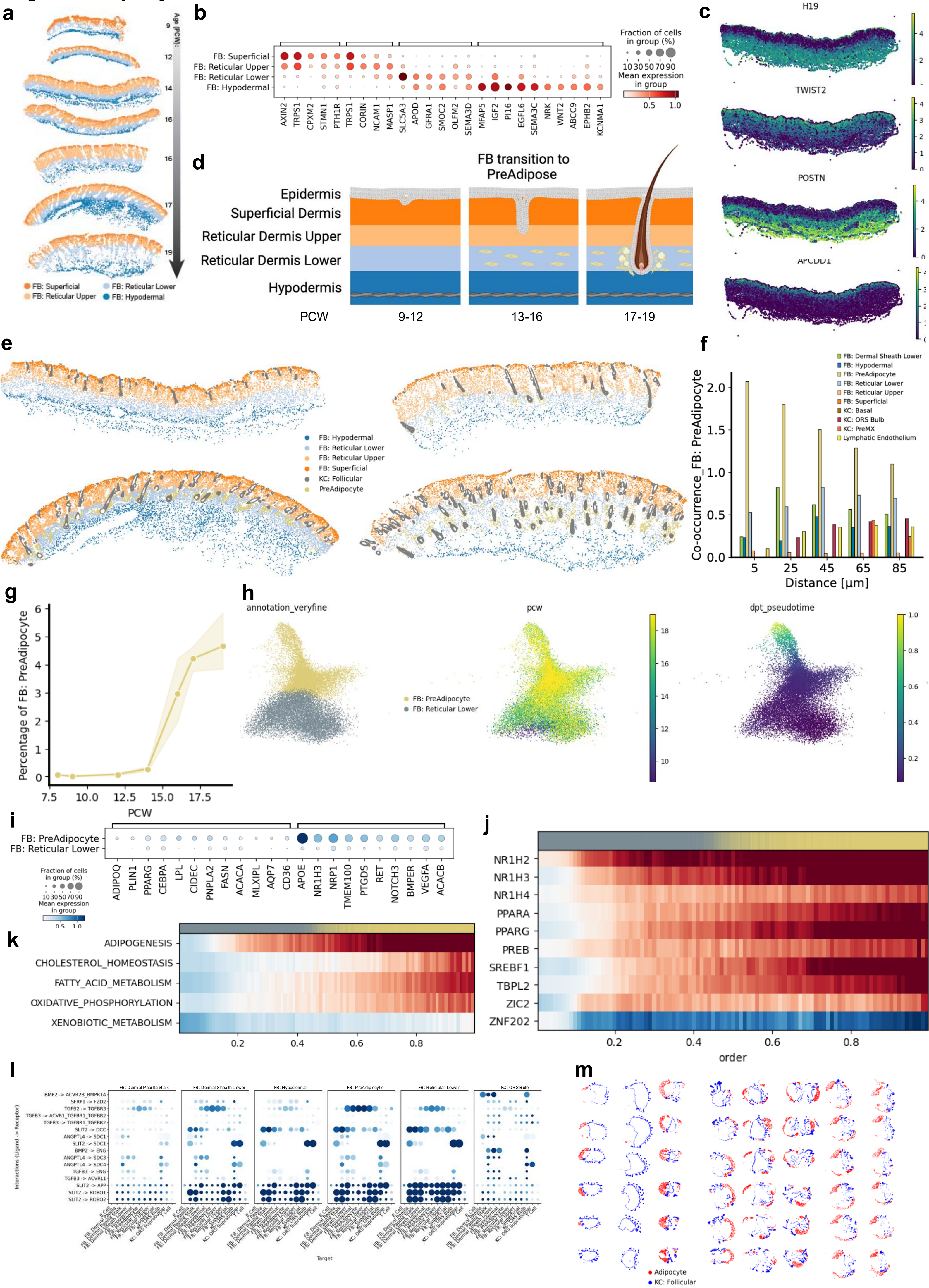

**Fig S8: Adipocytes form around hair bulb in lower reticular dermis**

a The Dermis can be stratified in 4 layers that are consistently found from PCW 8-19. The lowest layer refers to the developing hypodermis, while the middle two layers represent the lower and upper reticular dermis. The top layer is called Superficial dermis and refers to what we think later becomes the papillary dermis.

b Gene expression markers of different dermal fibroblast layers.

c Expression of gene with spatial gradient on a PCW14 skin section. *H19* and *POSTN* increase in expression the deeper the section while *APCDD1* and *TWIST2* decrease from uppermost to lowermost layer.

d During hair development, PreAdipocytes from on the lower reticular dermis, that then assemble around hair follicles.

e Spatial location of PreAdipocyte in fetal skin sections of increasing age. At PCW14 there are almost no PreAdipocytes, while in PCW19 they fully surround the bulbs of hair follicles.

f Co-occurrence of PreAdipocytes with dermal layers and other cell types at radii of 10 - 100 um. High co-occurrence can be observed between the PreAdipocytes and the lower reticular dermis as well the the lower dermal sheath that is wrapping around the bulb of hair follicles.

g Percentage of PreAdipocytes over time, shows strong increase of PreAdipocytes after PCW14.

h UMAP of reticular lower fibroblasts and PreAdipocytes coloured by annotation, real time (PCW) and Pseudotime.

i Differential gene expression between reticular lower fibroblasts and PreAdipocytes.

j TF activity along developmental trajectory from reticular lower fibroblasts to PreAdipocytes, showing high activity of fatty acid production TF like Liver-X-Receptor, PPARA/G and SREBF2.

k Hallmark pathway enrichment along the developmental trajectory from lower reticular dermis to PreAdipocytes constructed by computing a Pseudotime from Reticular Dermis to PreAdipocytes. Along this axis, we can see enrichment of adipogenesis, fatty acid metabolism and oxidative phosphorylation.

l LR based cell-cell communication analysis showing several potential interactions of dermal papilla stalk cells with lower dermal layers through Wnt-signals.

m Spatial location of Adipocytes (red) and follicular keratinocytes (blue) in the skin organoid.

### Fig S9: Companion layer forms consistently across scalp skin and skin organoid

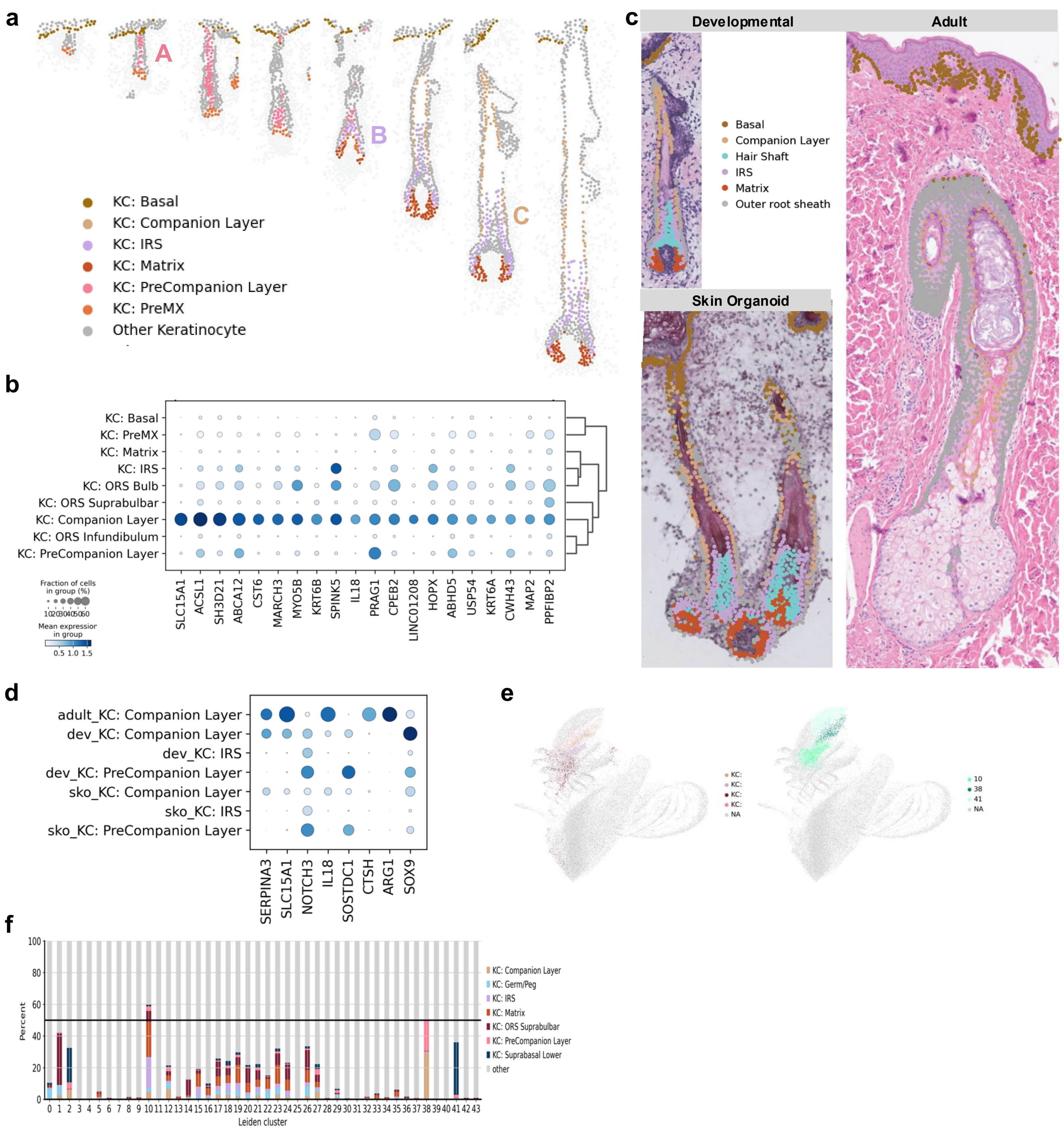


**Fig S9: Companion layer forms consistently across scalp skin and skin organoid**

a Cell type annotation showing the spatial location of matrix, inner root sheath and companion layer cells and progenitors. While PreMX and PreCompanion Layer are present from stage 4-7, stage 8-10 present the mature companion layer. Matrix and IRS however are present already one stage earlier at stage 7.

b Gene expression markers of developing companion layer.

c Spatial location of developmental, skin organoid and adult companion layer.

d Comparison of companion layer markers across development adult and skin organoid.

e UMAP of ATAC highly variable peaks. The spatial location shows proximity of companion layer cells with PreCompanion Layer.

f Percentage of Leiden clusters annotated as certain cell types. Cluster 38 show highest fraction of PreCompanion- and Companion Layer.

**Fig S10: PreCompanion Layer shows similarity to suprabasal keratinocytes**

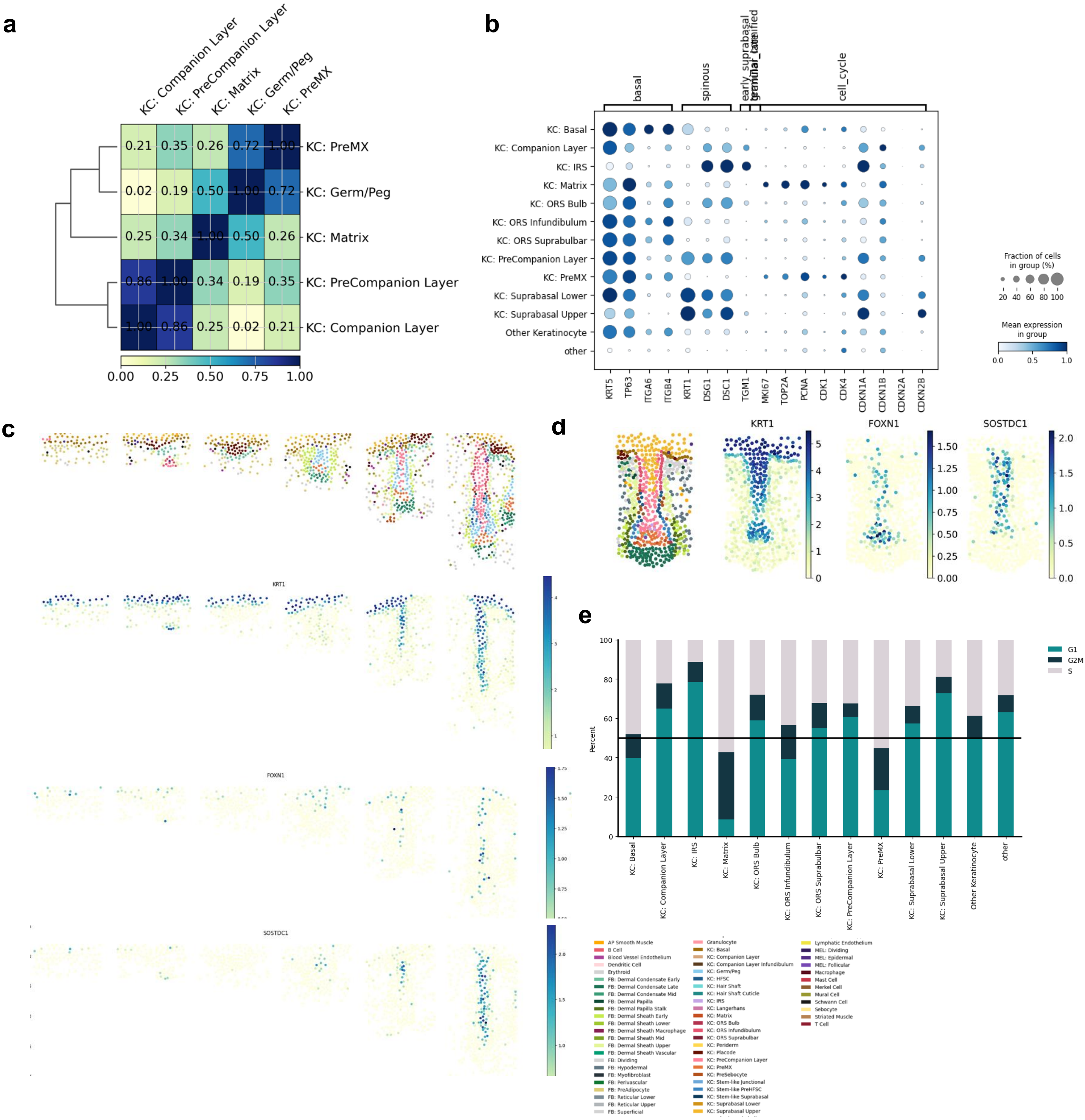


**Fig S10: PreCompanion Layer shows similarity to Suprabasal keratinocytes**

a Correlation heatmap of ATAC-peaks of follicular hair follicle populations.

b Dot plot showing cell cycle and keratinocyte maturation markers. PreCompanion layer cell share genes such as *KRT1, DSG1* and *DSC1* with the Suprabasal KCs. The latter two are also expressed in later ORS, IRS and Companion layer. PreCompanion layer and Suprabasal KCs also share expression of *CDKN2B*.

c Cell annotation and gene markers (*KRT1, FOXN1, SOSTDC1*) for PreCompanion layer in early-stage HF.

d Stage 6 Hair follicle from skin organoid displaying PreCompanion layer and KC Suprabasal in the middle of the Follicle.

e Cell cycle phases of follicular Keratinocytes.

### Fig S11: Inner root sheath and hair shaft upward movement is characterized by specific gene expression

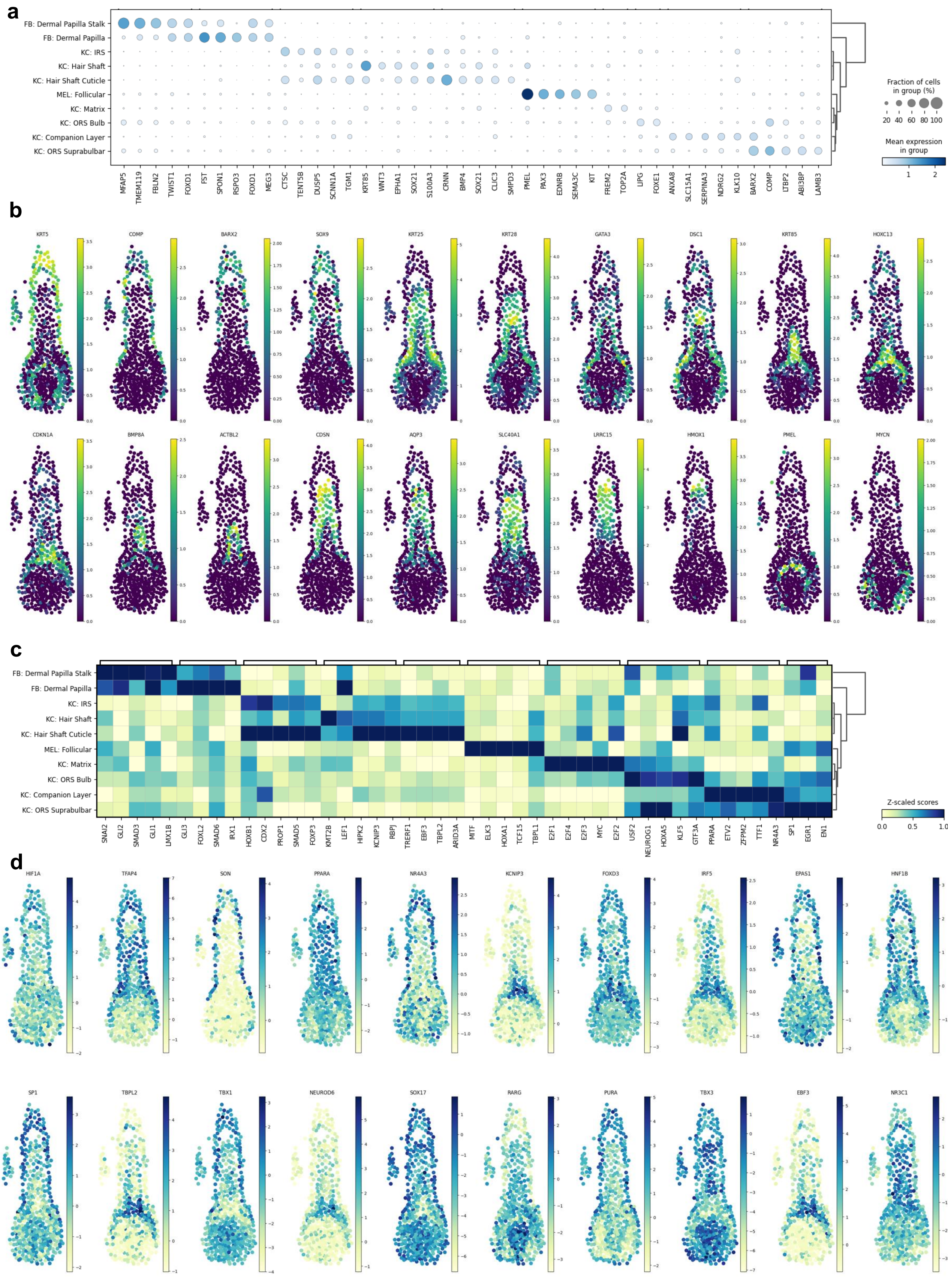

**Fig S11: Inner root sheath and hair shaft upward movement is characterized by specific gene expression**

a Dot plot of filtered gene expression markers of hair bulb relevant populations.

b Gene expression of spatially highly autocorrelated genes in a hair bulb section ordered after cell type of expression.

c Most specific TF activities in hair bulb relevant populations.

d Activity estimation of spatially highly autocorrelated TFs in a hair bulb section ordered after cell type of expression.

### Fig S12: IRS specification is driven by NOTCH activity consistent with skin organoid

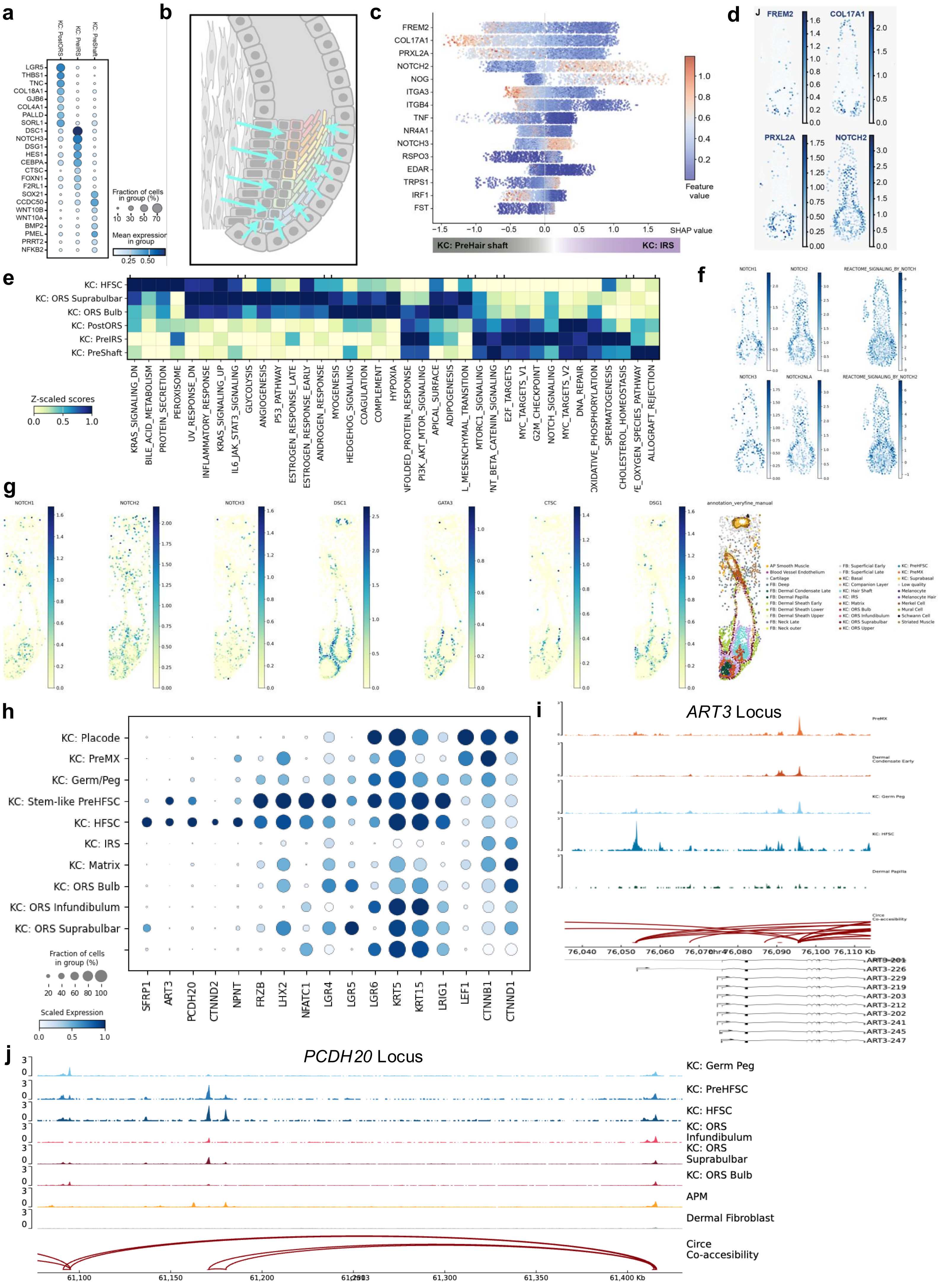

**Fig S12: IRS specification is driven by NOTCH activity consistent with skin organoid**

a Gene markers of matrix fine populations. While the PostORS still expression ORS markers like *LGR5*, the PreIRS strongly expresses IRS markers like *DSG1*, *CTSC* and *NOTCH3*. The PreShaft strongly expressed *WNT10A/B* as well as *SOX21*.

b Cartoon showing the location of the area where fate decisions between IRS and Hair Shaft are made in the developing bulb.

c ignite result for most predictive factor for PreIRS of PreShaft identity. While the top factors for the PreIRS *MFAP5* and *THY1* are related to notch signalling, the top PreShaft discriminators *FST* and *BMP7* are known signalling molecules expressed in the dermal papilla.

d Spatial location of top ignite factors: While *MFAP5* and *THY1* are expressed around the bulb in the dermal sheath, *FST* and *BMP7* are mostly expressed in the dermal papilla.

e Hallmark pathway enrichment showing high enrichment of notch signalling in PreIRS, while the Shaft population has the highest enrichment on WNT/Catenin signalling.

f NOTCH expression and Reactome enrichment show strong notch activity in IRS areas of the hair follicle.

g Skin organoid expression of NOTCH and IRS markers.

h Dot plot showing Bulge markers such as *SFRP1, ART3, PCDH20, CTNND1* and *NPNT*, generated from differential gene expression.

i ATAC-coverage around the ART3 locus, showing no accessibility of common transcription start site as well as high accessibility of alternative transcription start sire of ART3-226.

j ATAC-coverage around the PCDH20 locus, showing open chromatin 200-300 kb downstream of PCDH20.

### Fig S13: Lower ORS is characterized by stem cells markers and WNT-inhibition

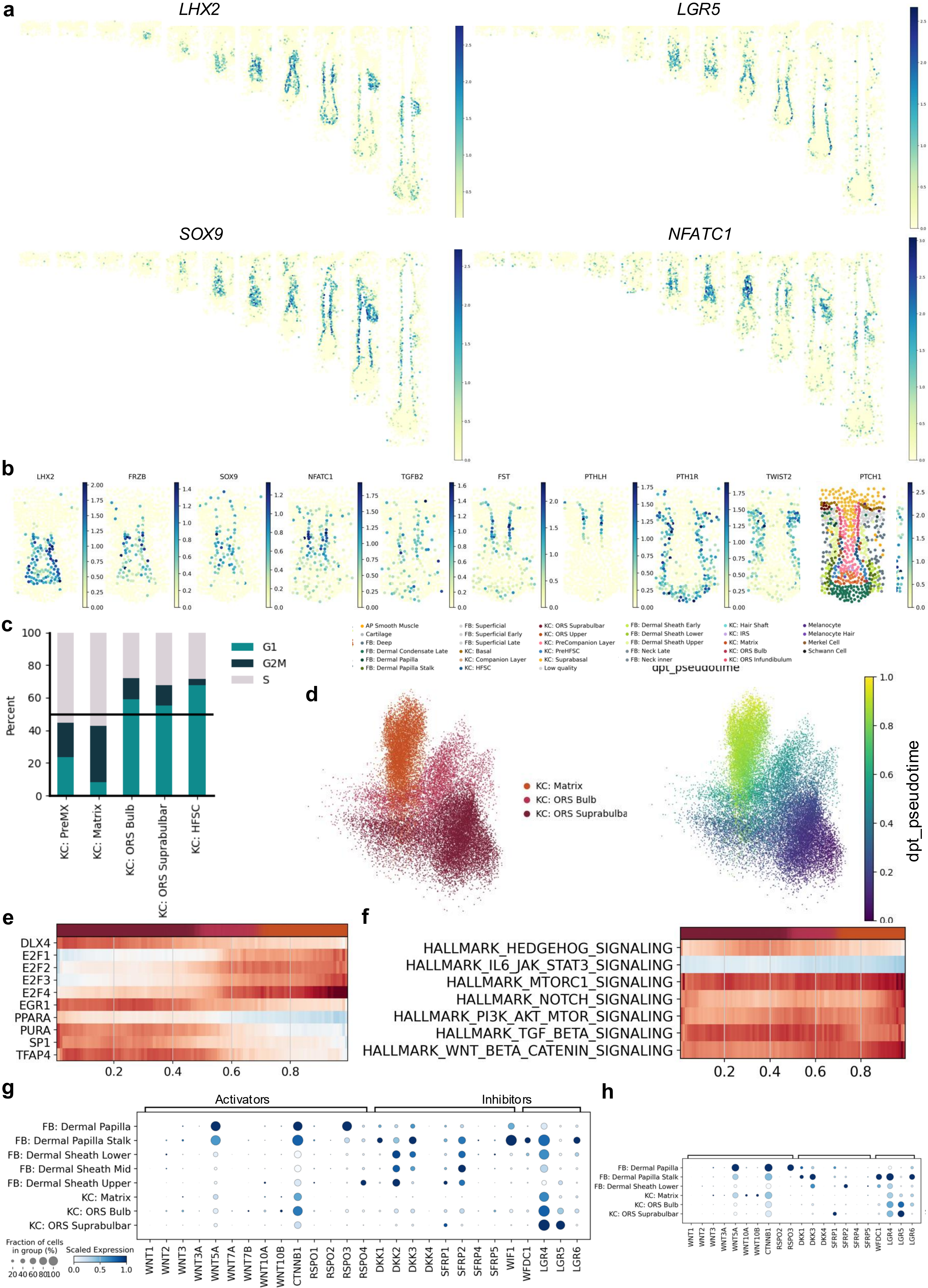

**Fig S13: Lower ORS is characterized by stem cells markers and WNT-inhibition**

a Expression of lower segment (*LHX2* and *LGR5*) and stemness markers (*SOX9* and *NFATC1*) in Stage 6 human hair follicles.

b Expression of segment and stemness markers in Stage 6 skin organoid hair follicles.

c Cell cycle phases per cell type. Matrix cells have a very high rate of proliferation rate measure by a very low percentage of cells in G1 phase. PreMX is also proliferative but less than Matrix. ORS and HFSC have lower proliferation rates with more then 50 percent in G1.

d UMAP showing the cell type annotation and Pseudotime of ORS suprabulbar, bulbar and KC: Matrix.

e TF activity along Pseudotime show strong increase in E2F activity upon entering late bulb/matrix stages.

f MYC upstream pathway activity along pseudotime, a constant increase in activity can be observed for WNT and less so for NOTCH signalling.

g Single-nuc gene expression of WNT activators, inhibitors and modulators show specific expression of wnt inhibitors and modulators in dermal papilla stalk and strong expression of RSPO3 in dermal papilla.

h Skin organoid expression of WNT activators, inhibitors and modulators .

### Fig S14: Arrector pili muscle develops smooth muscle phenotype

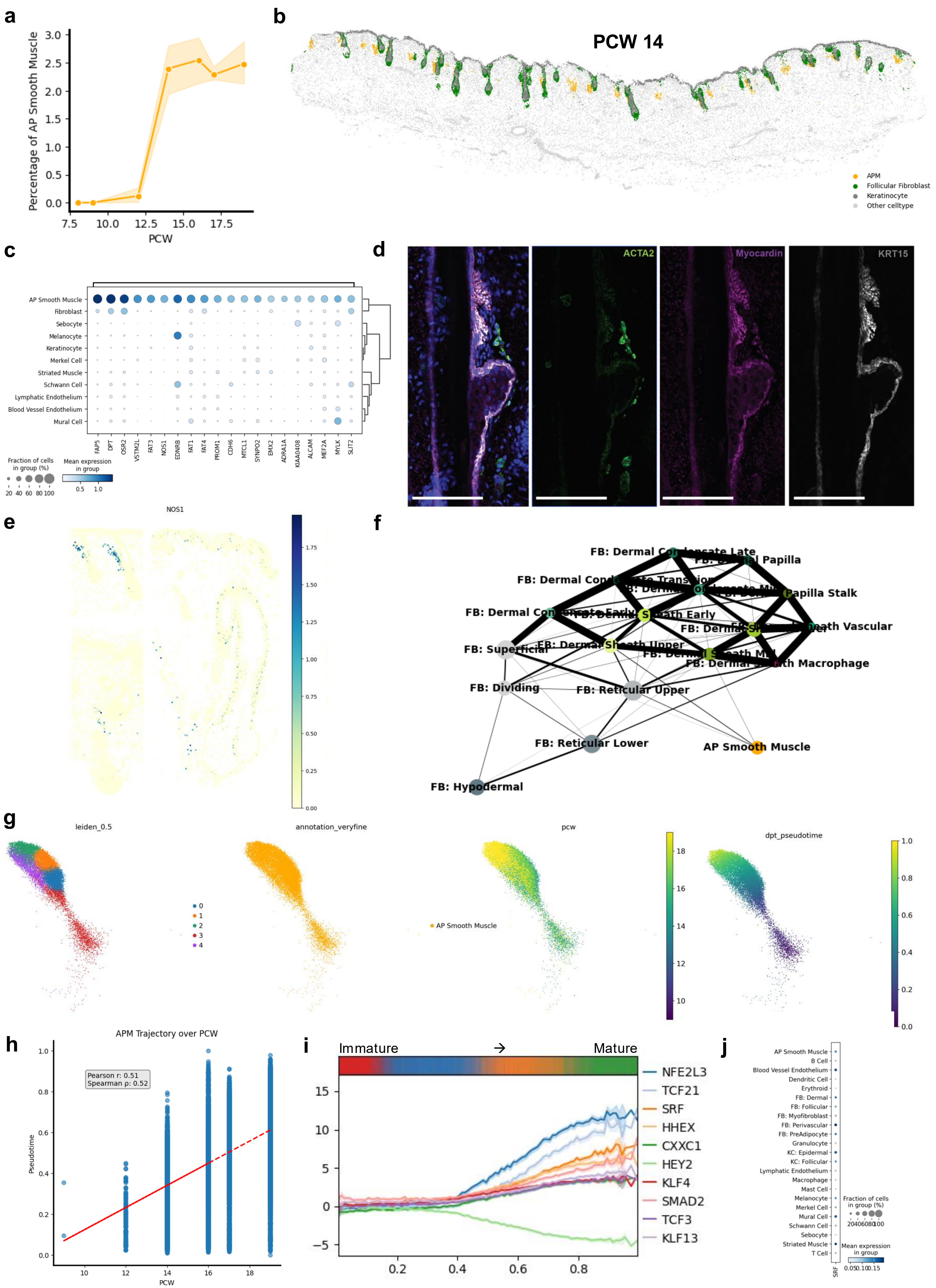

**Fig S14: Arrector pili maturation is driven by Myocardin activity**

a Per section percentage of APM cells over time (PCW). APM cell abundance increases after PCW12.

b Spatial location of APM cells (yellow) at PCW 14, next to hair follicles surrounded by follicular fibroblasts (green).

c Xenium gene expression markers of the APM cells.

d Immunohistochemistry of arrector pili muscle stained by ACTA2 and Myocardin.

e Spatial location of NOS1 expression in developmental, SkO and adult arrector pili muscles.

f PAGA plot describing transcriptional similarity with different dermal (grey) and follicular (green) fibroblast populations.

g UMAP of APM cells coloured by Leiden clustering (left), cell type annotation (middle left), real time in PCW (middle right) and Pseudotime (right).

h Correlation of Pseudotime with actual time r = 0.51, p =0.52.

i Line plot showing the TF activity along Pseudotime shows diversification of transcriptional activity after roughly the first third of the axis.

j Dot plot showing SRF expression across all cell types.

### Fig S15: Arrector pili maturation is driven by Myocardin activity

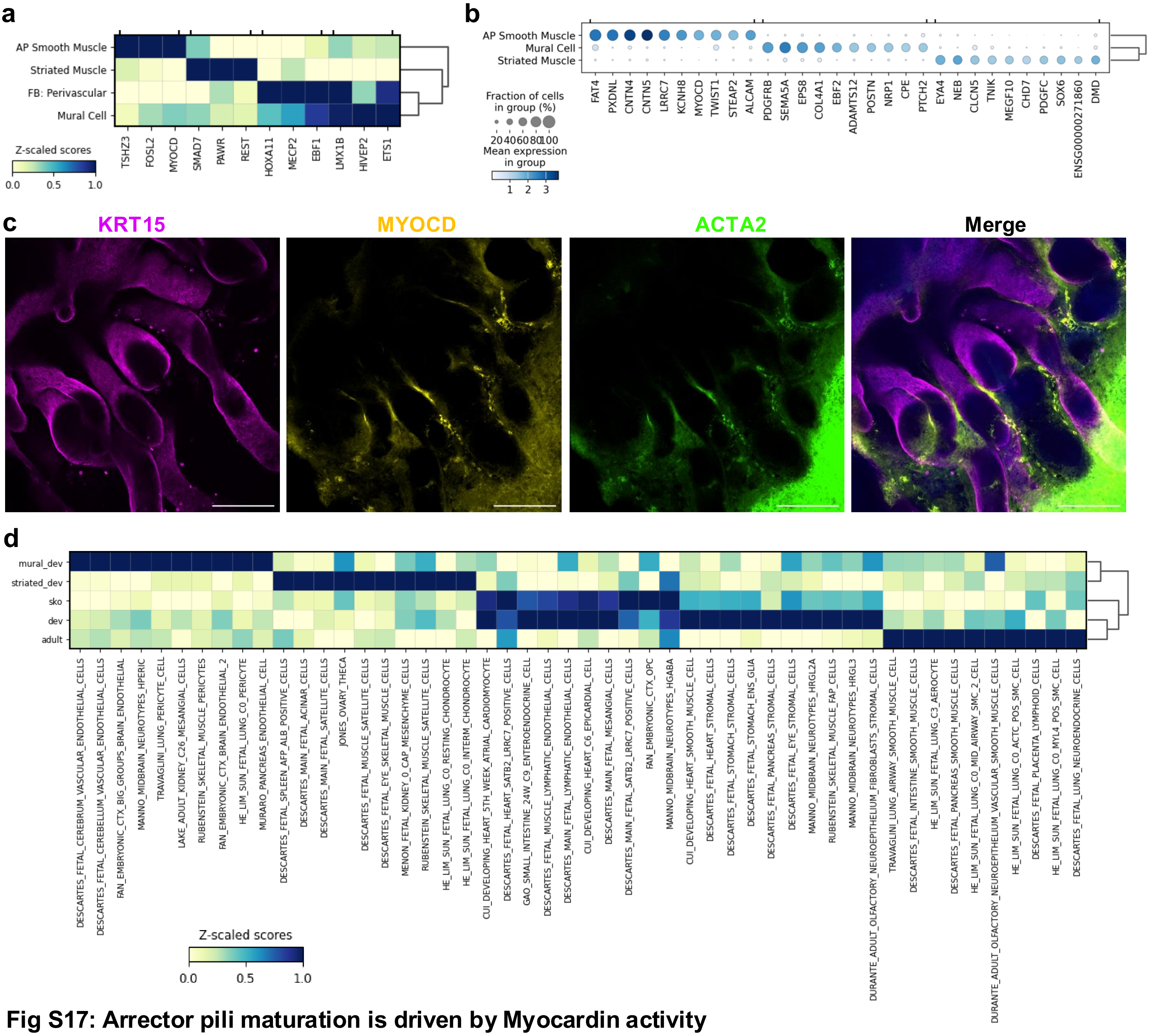


**Fig S17: Arrector pili maturation is driven by Myocardin activity**

a Matrix plot showing the TF activity among different muscle associated.

b Dot plot showing the main markers of muscle cells in the skin organoid.

c Whole mount of 145-day old skin organoid stained by *KRT15, MYOCD* and *ACTA2*.

d Matrix plot showing cell type signature enrichment among developmental muscle, skin organoid and adult APM both skin organoid and development have high enrichment of fetal heart and cardiomyocyte signatures.

### Fig S16: Developing sebaceous gland shows lipid accumulation

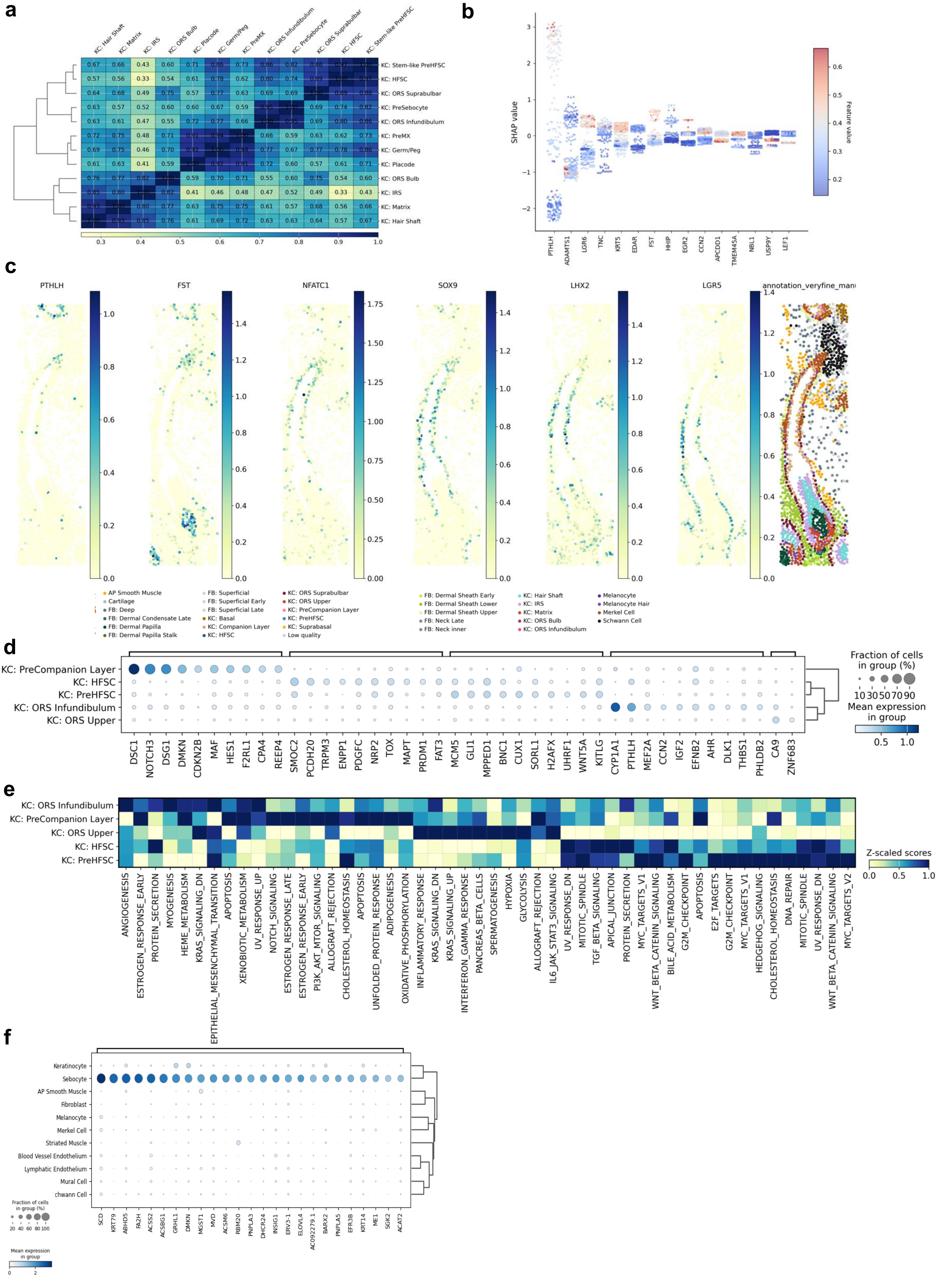

**Fig S16: Developing sebaceous gland shows lipid accumulation**

a Transcriptional correlation of ORS keratinocytes show strong similarity of PreSebocytes with KC: ORS Infundibulum and KC: ORS PreHFSC to KC: ORS Suprabulbar

b Ignite prediction factor for environmental factors that drive PreSebocyte vs KC: ORS Infundibulum (model roc_auc: 0.84) .

c Expression of segment and stemness markers in hair follicle of mature skin organoids.

d Skin organoid gene expression markers of follicular keratinocytes.

e Hallmark pathway enrichment of skin organoid follicular keratinocytes.

f Sebocyte marker genes in the single-nuc RNA dataset show Sebocyte specific expression of many fat and cholesterol production related genes.

### Fig S17: Sebaceous gland development is characterized by p21 driven cell cycle inhibition

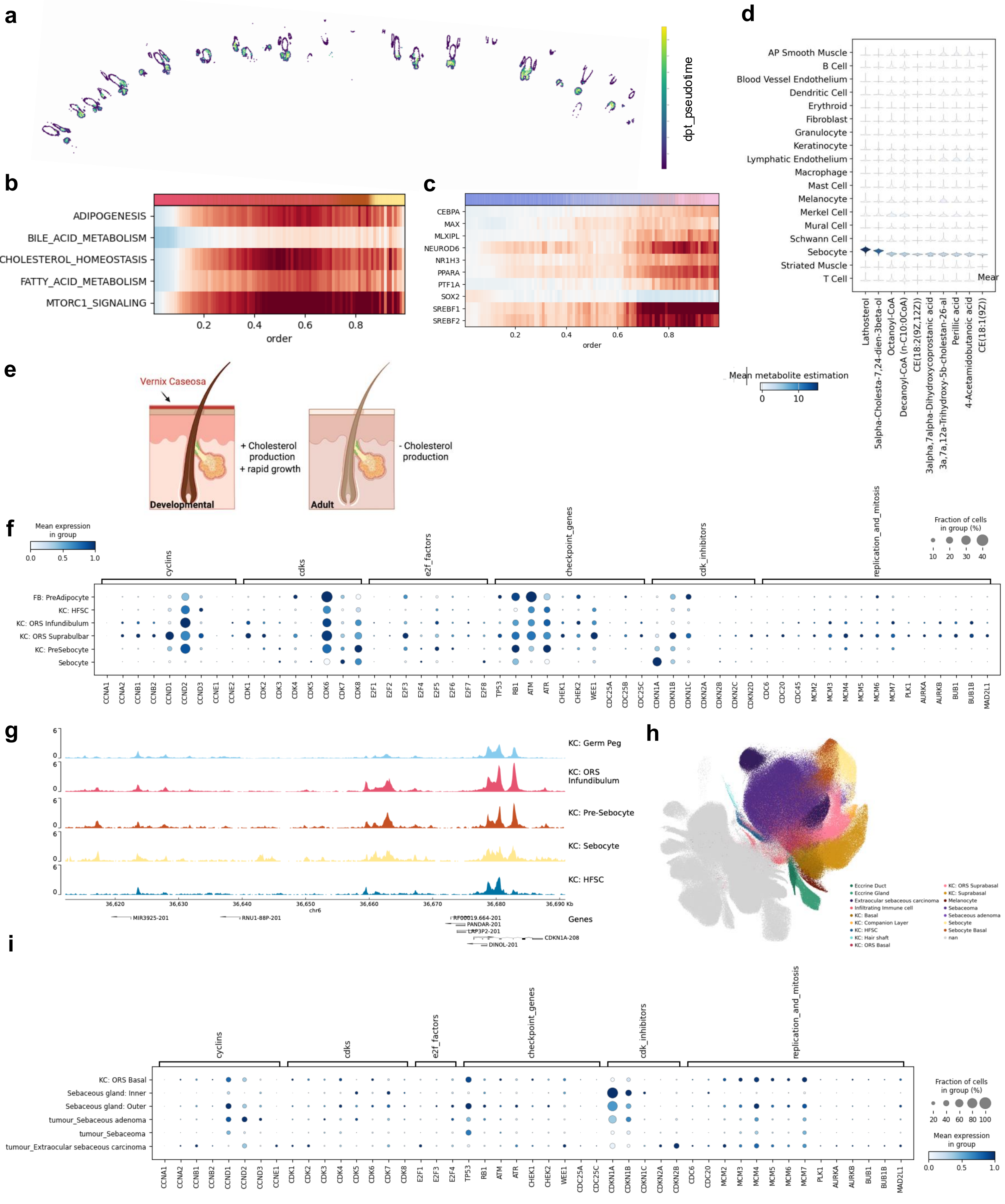

**Fig S17: Sebaceous gland development is characterized by p21 driven cell cycle inhibition**

a Spatial location of sebocyte developmental program (Pseudotime) values shows association with location towards the center of the gland.

b Top pathways associated with sebocyte developmental program show lipid associated metabolism and signalling.

c Adult sebocyte trajectory shows similar activation of TF to development.

d Enrichment of metabolic enzymes using MetalinksDB. Sebocytes show high enrichment of Lathosterol related enzymes.

e Cartoon showing key differences between adult and developmental sebocytes that include cholesterol production and the vernix caseosa.

f Expression of cell cycle related genes in sebocytes and sebocyte related cells, showing that CDKN1A is the only cell cycle inhibitor highly expressed and specific to sebocytes .

g Upstream region of CDKN1A showing several regions opening in PreSebocytes and sebocytes that can be putative regulatory sites.

h UMAP showing the annotation of the Sebaceous tumor dataset.

i Gene expression of p21 (CDKN1A), across tumor and non-tumor cells.
